# Adaptive Baseline Fitting for ^1^H MR Spectroscopy Analysis

**DOI:** 10.1101/2020.02.17.949495

**Authors:** Martin Wilson

**Affiliations:** Centre for Human Brain Health and School of Psychology, University of Birmingham, Birmingham, UK

**Author notes:** Correspondence Martin Wilson, Centre for Human Brain Health, University of Birmingham, Edgbaston, Birmingham, B15 2TT, United Kingdom.

**Keywords:** ABfit, MRSI, spectral analysis, automated, open-source, spline

## Abstract

**Purpose:** Accurate baseline modeling is essential for reliable MRS analysis and interpretation — particularly at short echo-times, where enhanced metabolite information coincides with elevated baseline interference. The degree of baseline smoothness is a key analysis parameter for metabolite estimation, and in this study a new method is presented to estimate its optimal value.

**Methods:** An adaptive baseline fitting algorithm (ABfit) is described, incorporating a spline basis into a frequency-domain analysis model, with a penalty parameter to enforce baseline smoothness. A series of candidate analyses are performed over a range of smoothness penalties, as part of a four stage algorithm, and the Akaike information criterion is used to estimate the appropriate penalty. ABfit is applied to a set of simulated spectra with differing baseline features and experimentally acquired 2D MRSI — both at a field strength of 3 Tesla.

**Results:** Simulated analyses demonstrate metabolite errors result from two main sources: bias from an inflexible baseline (underfitting) and increased variance from an overly flexible baseline (over-fitting). In the case of an ideal flat baseline ABfit is shown to correctly estimate a highly rigid baseline, and for more realistic spectra a reasonable compromise between bias and variance is found. Analysis of experimentally acquired data demonstrates good agreement with known correlations between metabolite ratios and the contributing volumes of gray and white matter tissue.

**Conclusion:** ABfit has been shown to perform accurate baseline estimation and is suitable for fully-automated routine MRS analysis.

## 1 INTRODUCTION

A number of key metabolites may be detected using ^1^H Magnetic Resonance Spectroscopy (MRS), providing a non-invasive measure of healthy and diseased brain tissue metabolism. Clinical applications include the assessment of brain tumors, metabolic disorders and neonatal encephalopathy [1, 2] where the concentration of certain metabolites may inform disease diagnosis or predict patient outcome. Further applications are present in the neuroscience and psychiatry domains, with particular interest in the direct detection of neurotransmitter levels such as GABA and glutamate — which have been shown to be abnormal in Schizophrenia [3] and modulate in response to tasks [4, 5].

MRS scans are typically performed at short (30 ms) or long (144 ms) TE’s, with short-TE scans being preferred due to reduced T2 relaxation and dephasing of multiplets resulting in improved metabolite detection sensitivity [6]. However, short-TE scans are typically more susceptible to artifacts originating from insufficient water and scalp lipid suppression, in addition, broad signals from macromolecules also become enhanced [7]. Residual water signals, lipid signals and macromolecules all have the potential to bias metabolite measurements due to spectral overlap and interference. Therefore, appropriate analysis methodology is particularly important to achieve the full benefit of MRS at short-TE.

Parametric fitting is currently the most widely used analysis method, and typically incorporates a set of simulated or experimentally measured metabolite and macromolecule signals — known as a basis set. An important distinction between analysis methods is their approach for mitigating metabolite estimation bias from broad signals not present in the basis set, usually referred to as “baseline modeling”. The true baseline should only contain artifacts, such as extraneous lipid or water signals, however baseline modeling inaccuracy results in signals being fully or partially attributed to the wrong source. One of the most popular baseline modeling methods incorporates a set of smooth spline functions into the fitting procedure, with additional smoothness imposed by penalizing greater baseline complexity. The LCModel [8] and AQSES [9] algorithms both use penalized spline baseline modeling, with analysis performed in the frequency-domain and time-domain respectively.

An alternative approach to baseline modeling exploits the rapid decay of baseline signals in the time-domain by omitting the preliminary data points during the fitting process, reducing their interference with the more slowly decaying metabolites. The QUEST [10] and TARQUIN [11] methods both use this time-domain truncation approach. The FITT [12] algorithm combines the wavelet transform with Lowess filtering in the frequency-domain to separate metabolite and baseline signals. In addition to metabolite signals, it is often beneficial to add known lipid and macro-molecular signals to the basis set — particularly for the analysis of short-TE MRS [13]. At the time of writing, the LCModel and TARQUIN algorithms include a set of these signals by default, whereas QUEST, AQSES and FITT require these signals to be manually appended to the metabolite basis.

Control over the level of baseline flexibility (or smoothness) is a common and necessary requirement of each of the baseline modeling methods outlined above. In spline based approaches, a combination of the number of spline functions for a given frequency range and the smoothness penalty parameter control the baseline flexibility. For LCModel, the frequency spacing between the spline basis functions is dependent on data quality, and is set at maximum of 1.5 times the estimated full width at half maximum (FWHM) of the metabolite resonances or 0.1 ppm [8]. Similarly, for the FITT algorithm a fixed Lowess filter smoothing value is used and wavelet coefficients with scales less than twice the FWHM are excluded from the baseline model to ensure smoothness [12]. In the time-domain truncation approach baseline flexibility is primarily determined by the number of initial data points to be omitted from the fit evaluation. For QUEST and TARQUIN the number of truncated data points, and therefore degree of baseline flexibility, is set at a default value that may be adjusted by the user.

Automated methods to determine the correct degree of baseline flexibility are important for obtaining accurate metabolite levels independently of the analyst. Furthermore, the manual adjustment of baseline flexibility for each individual spectrum is impractical for MRSI studies — where hundreds of spectra may be acquired in a single scan. Whilst LCModel provides automated adjustment of baseline flexibility, a growing number of analysts choose to manually override the default analysis settings by adjusting the spline spacing parameter (DKNTMN). The first reported use of this manual adjustment was to improve the modeling of macromolecular resonances in rat brain at 9.4 T [14]. More recently, this parameter has been adjusted to encourage flatter baselines [15, 16, 17], suggesting the default LCModel baseline flexibility may not be optimal in some cases.

Finding the optimal degree of baseline flexibility is a crucial question in MRS analysis research, yet few studies have investigated this topic in detail. Using simulated data, Ratiney et al. demonstrated how the interference between metabolite and baseline signals was reduced by increasing the number of omitted data points, but this came at the cost of inflating errors due to noise [18]. More recently, Near et al. showed how the estimated baseline in LCModel can depend strongly on spectral SNR and metabolite FWHM, and that errors caused by baseline instability may dominate over errors from spectral noise in some cases [19]. The influence of baseline flexibility has also been explored using experimentally acquired data, with a recent study demonstrating a 15% difference in metabolite levels when comparing between a default and less flexible baseline model [20]. Baseline flexibility has also been shown to have a strong influence of the measurement of 2-hydroxyglutarate [21].

In this study, we introduce a new method to automatically determine the optimal degree of baseline flexibility for a frequency-domain spline based fitting algorithm. Firstly, background is given on the use of penalized splines for optimal data smoothing. A fully-automated fitting algorithm is presented, incorporating a novel method to automatically estimate the optimal level of baseline flexibility. Finally, the new method is validated with simulated and experimentally acquired MRSI data.

## 2 THEORY

### 2.1 Penalized spline smoothing

Baseline signals have a characteristically smooth spectral appearance, and must be accurately modeled to avoid biasing metabolite estimates. In good quality ^1^H MRS of brain tissue baseline signals have a low intensity, relative to the primary metabolite resonances, and are therefore challenging to estimate in the presence of noise. Estimating a smooth function from noisy data is known in statistics as “scatterplot smoothing”, and a number of approaches have been developed [22]. In this section we briefly outline the method of penalized splines in the simpler context of scatterplot smoothing, before describing their use as part of an MRS fitting algorithm.

A spline is a piecewise function made up of one or more polynomial segments joined together at points known as “knots”. A wide range of spline functions are possible, however Basis-splines, more commonly known as B-splines, are a popular choice for smoothing applications due to their favorable numerical properties [23]. The degree of a B-spline function determines its overall smoothness, and third degree (or cubic) B-splines are often used for smoothing. A cubic B-spline basis is show in Figure 1A with an offset added in the y-axis for clarity. B-spline bases consist of regularly spaced overlapping spline functions, spanning the full range of x values.

**FIGURE 1.**
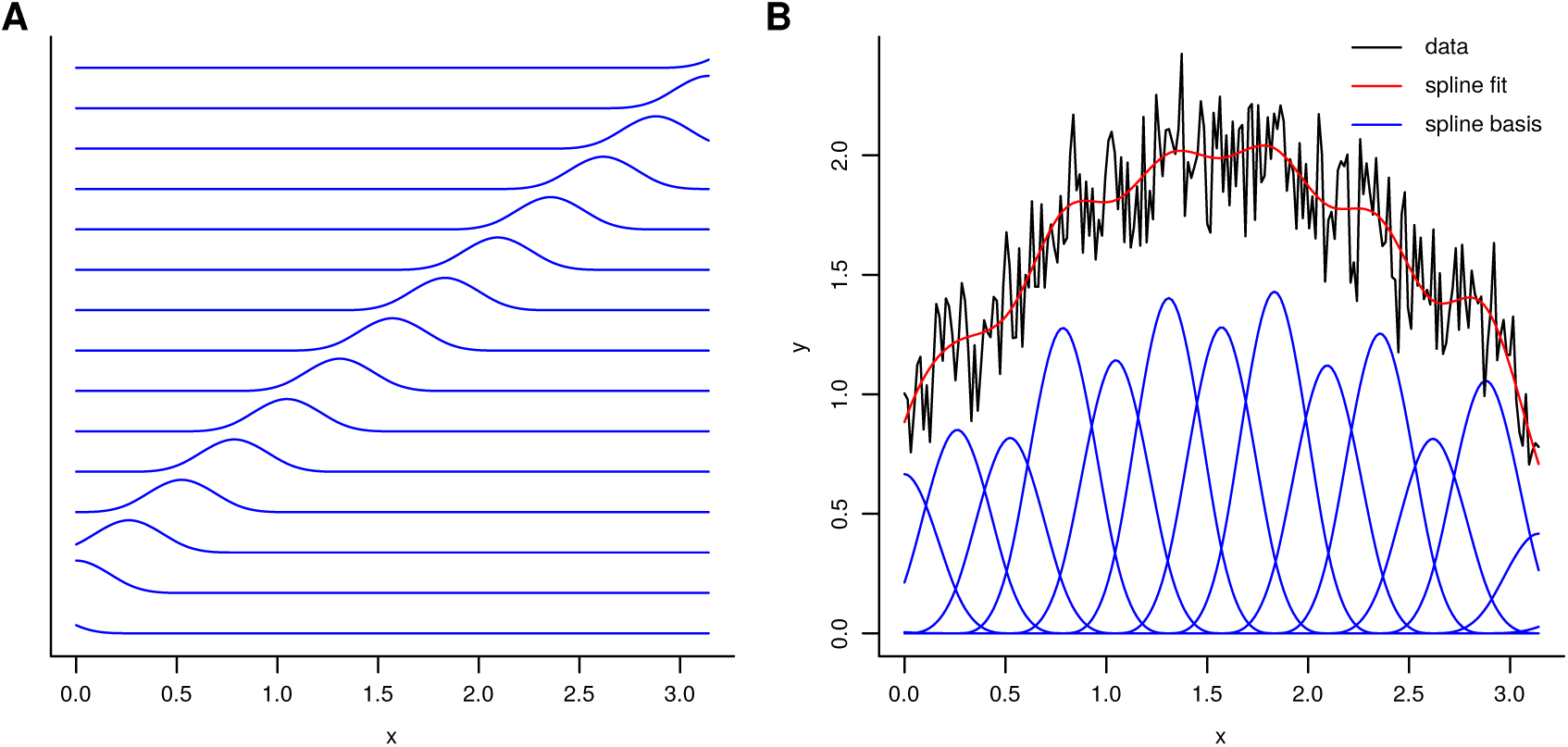
A) Cubic B-spline basis of 15 components with each component offset vertically to reduce overlap. B) P-spline regression of a noisy signal with an underlying smooth trend.

A B-spline basis may be used to obtain a smooth estimate of a signal using simple linear regression. Figure 1B shows the result of a spline regression, where each spline function has been optimally weighted, such that the sum of the functions (spline fit line) is the least squares fit to the data. The desired smoothness of the fit is controlled by adjusting the spline knot spacing, which in turn changes the density of spline functions. In the case of Figure 1B, 15 functions were found to give a reasonable level of smoothness. Increasing the density of spline functions allows more detail to be captured by the fit, however too many functions results in an increased sensitivity to noise and the smooth estimate begins to exhibit random fluctuations. Conversely, insufficient density of spline functions results in the spline fit being unable to model genuine smooth trends present in the data.

An alternative to adjusting the number of spline basis functions to achieve a desired level of smoothness is to introduce a penalty parameter into to spline regression model. The smooth estimate **ŷ**, of our data **y**, is calculated as: **ŷ** = **B â**, where **B** is a B-spline basis in matrix form, and **â** is a vector of corresponding spline weightings to be determined. In simple spline regression **â** is found by solving the normal equations to minimize the sum of the squared differences between **y** and **ŷ**. In penalized spline regression, the minimization function *Q*_*B*_ is adjusted to incorporate an additional term to enforce smoothness in the estimate:

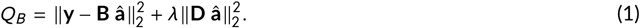

The *λ* parameter controls the degree of smoothness by penalizing solutions for **â** that interact with the difference matrix **D**. The difference matrix may be constructed to penalize the first, second or higher orders of differences between **â** values. Here, we exclusively use a second order difference matrix, which is particularly effective for modeling the smooth baseline features typically found in MRS data.

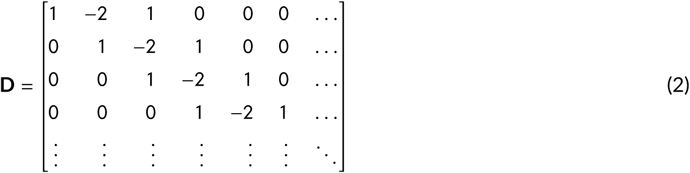

An example of the second order difference matrix is given in (2), which shows how increased differences between adjacent weighting factors in **â** consequently increase the penalty term in Equation (1). The minimization function *Q*_*B*_ represents a compromise between minimizing the fit residual and smoothness, where a value of zero for *λ* results in simple spline regression. Larger values of *λ* encourage a smoother **ŷ**, to the point where **ŷ** becomes a straight line fit for very large penalties when using a second order difference matrix. Approximately linear baselines, encouraged by a second order difference matrix, act as a good model for the tails of the residual water resonance typically found in MRS data. The solution to Equation (1) may be found by row-wise concatenation (augmentation) of **B** and 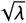 **D**, and appending zeros to **y** to match the number of combined rows:

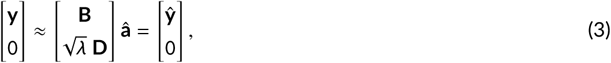

and regressing the augmented **y** vector on the augmented **B** matrix to yield **â**.

The general approach for penalized spline regression is to over-specify the number of B-spline basis functions, and primarily control the smoothness through the adjustment of *λ* acting on a difference matrix. We refer to this approach as “P-splines”, first introduced by Eilers and Marx [24]. Whilst the value of *λ* has a direct effect on smoothness, it is an unintuitive parameter to interpret, as the optimal value often varies by several orders of magnitude. In addition, *λ* has a complex dependence on the density of the spline functions, the number of data points and other factors unrelated to smoothness. A more intuitive measure of the smoothness of a P-spline model is known as the effective dimension (ED), proposed by Hastie and Tibshirani [25]. For a given value of *λ*, B-spline basis **B** and difference matrix **D**, we calculate ED as follows:

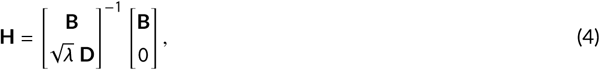

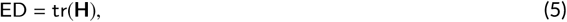

where tr denotes the trace of a matrix. For a small value of *λ*, ED approaches the number of spline functions in the basis **B**, and for large values, ED approaches 2 when using a second order difference matrix since a straight line fit has two degrees of freedom: the gradient and the y-intercept. Similarly, a heavily penalized first order difference matrix approaches an ED of 1, resulting in a perfectly horizontal baseline with one degree of freedom corresponding to the y-intercept value. Note, in contrast to the LCModel parameter for adjusting baseline flexibility (DKNTMN), a smaller value of ED corresponds to a smoother baseline estimate.

Using simulated data we investigate the relationship between *λ*, ED and the optimal level of smoothness. The top left panel in Figure 2A shows a simple sine function with added normally distributed noise, shown in red and black respectively. Candidate P-spline smoothers, with differing values of *λ*, are shown in the remaining 5 panels. Since the true shape of the underlying smooth function is known, the error of the P-spline estimate may be calculated as the sum of squared differences between the true function and the estimate. A plot of the error as a function of *λ* is shown in Figure 2B. The sum of squared differences between the P-spline estimate and noisy data (residual) and the ED are also shown as a function of *λ* in part B.

**FIGURE 2.**
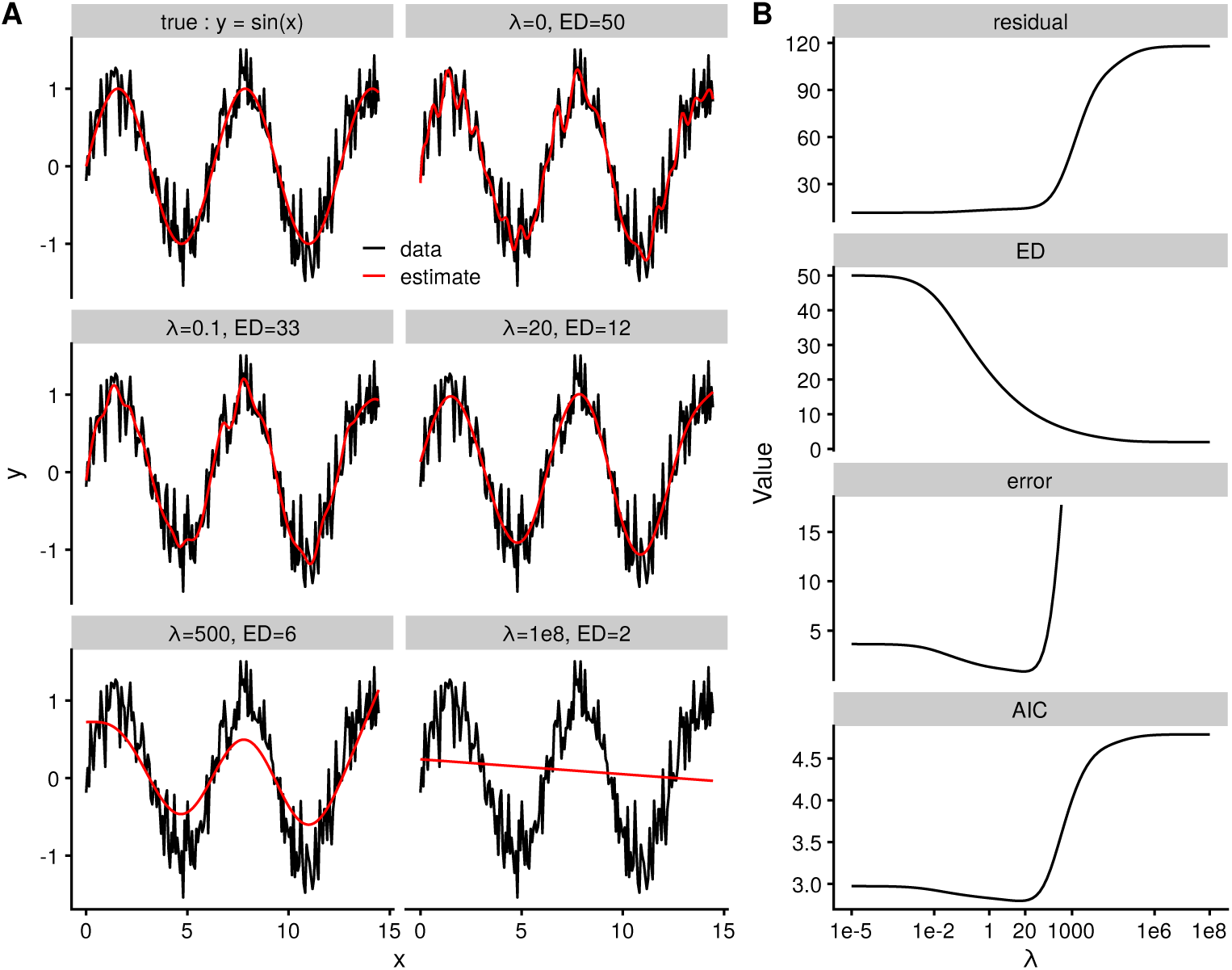
A) Penalized spline smoother applied to a simulated sine function over a range of smoothness penalties *λ*. B) Plots of the fit residual, smoother effective dimension (ED), fit error and Akaike information criterion (AIC) as a function of *λ*.

Inspection of the error plot reveals the optimal *λ* to be approximately 20 — corresponding to an ED value of 12, and this can be intuitively verified from part A. For larger values of *λ*, the estimate approaches a straight line fit — failing to capture the details in the sine cure and resulting in an increasing fit residual and fit error (part B). Models with insufficient freedom to adapt to genuine trends in the data result in biased estimates and this is known as “underfitting”. Conversely, too much freedom in the smoothing model (small *λ*) results in the estimate becoming overly sensitive to random fluctuations in the data resulting in “overfitting”. In least-squares fitting, there is a temptation to equate the smallest residual with the optimal fit, however Figure 2B clearly shows an increase in the error for *λ* values of less than 20 — despite a steady reduction in the residual.

Determining the optimal value for the smoothness parameter is one of the primary challenges for P-spline smoothing. A careful balance between instability from overfitting, and bias from underfitting, must be made for the most accurate estimate, and searching for the smallest fit residual is a useful, but insufficient metric of quality. Numerous approaches have been proposed to find the optimal smoothing value [22], and in this study we use *Akaike’s information criterion* (AIC) [26]:

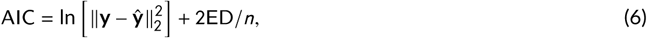

where ln denotes the natural logarithm and *n* is the number of data points. The AIC is typically used to compare models, with lower values indicating an improved balance between bias and instability. A commonly used alternative is known as the *Bayesian information criterion* (BIC), however the AIC is chosen here for its simpler form, which may be intuativly modified — as shown in the following section. A plot of the AIC as a function of *λ* is shown in the lower panel of Figure 2B, and the *λ* value with the lowest AIC shows good agreement with the true minimum error.

### 2.2 MRS baseline estimation using P-splines

In ^1^H MRS analysis, the baseline signal must be estimated in the presence of numerous overlapping metabolite lipid and macromolecule signals. Fortunately, these non-baseline signals have a known molecular origin and are therefore well characterized and accurately simulated from established parameters [27]. We can update Equation 3 to incorporate these additional molecular components by arranging into columns of the basis matrix **M**, and appending to the P-spline basis **B**:

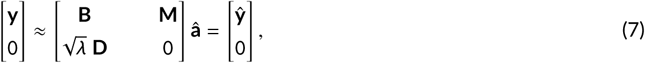

As with Equation 3, we regress the basis matrix on **y** to yield **â**, however since metabolite signal amplitudes are always positive, analysis stability may be improved by enforcing a non-negative constraint on the subset of **â** values corresponding to the weightings on the basis set **M** (**â**_*M*_ ≥ 0). The active-set method developed by Lawson and Hanson [28] is used herein to find the least-squares solution under the non-negative constraint — while still allowing the **â** values corresponding to the spline basis (**â**_*B*_) to remain unconstrained. Note that starting values are not required for any of the metabolite, macromolecular, lipid or spline basis function amplitudes when using this method.

A simulation study was performed to investigate the relationship between baseline smoothness, the accuracy of metabolite estimates and the AIC. Metabolite signals were simulated from known parameters [27] at levels consistent with normal brain tissue [29] — listed in Supporting Information Table S1. An experimentally derived macromolecule profile was also included in the simulation and basis matrix to yield a realistic spectrum [30]. This profile was comprised of multiple broad resonances combined with fixed relative intensities, and corresponded to only one element in the basis set. Simulated experimental conditions consisted of a field strength of 3 T, and semi-LASER localization (TE=28 ms). 1024 complex points were generated at a sampling frequency of 2000 Hz, 6 Hz Gaussian line-broadening was applied prior to zero-filling to 2048 points and Fourier transform to the frequency-domain. Ideal pulses were assumed and relaxation effects were not considered.

Ideal spectra were distorted with normally distributed noise resulting in a spectral SNR of 54 — typical for ^1^H MRS acquired from the human brain at 3 T. Signal strength was measured as the highest data point from the nominal NAA singlet peak at 2.01 ppm. A broad Gaussian resonance was added at 1.3 ppm with a linewidth of 100 Hz to simulate baseline distortion originating from scalp lipids. The degree of P-spline baseline flexibility is defined as the baseline ED (effective dimension) per ppm, which is more easily compared across analyses due to its independence from the number of points in the fit and the number of spline basis functions used. For example, an ED per ppm of 5 corresponds to an ED value of 19 when fitting the spectral region between 4 and 0.2 ppm (5 × (4 − 0.2)). Metabolite level estimates (**â**) were calculated using Equation 7 from real-valued data points in spectral region between 0.2 and 4 ppm, over a range of 10 levels of baseline flexibility. 32 spectra were analyzed at each level of baseline flexibility, with only the noise samples – drawn from the same distribution – randomly differing between each spectrum. The metabolite estimation error was calculated from the sum of the squared differences between the true amplitudes, listed in Table S1 (**a**), and estimated values (**â**) for each spectrum. Errors in the estimation of macromolecular or lipid signal amplitudes in the basis set were not considered. The mean and standard deviation of metabolite estimation errors was calculated across the 32 spectra at each level of baseline flexibility to evaluate accuracy and consistency. Error values are provided in absolute units as further scaling was not performed.

Metabolite estimation errors are shown in Figure 3A, displaying a comparable shape to the simpler model in Figure 2B. Over the range of baseline flexibility studied, underfitting with an inflexible baseline results in much greater metabolite estimate errors compared to overfitting — which can be verified by inspecting the fit result plots in Figure 3 parts C-F. Good agreement is also seen between the metabolite estimate error and the AIC curve (part B), with a low AIC value corresponding to the most accurate level of baseline flexibility.

**FIGURE 3.**
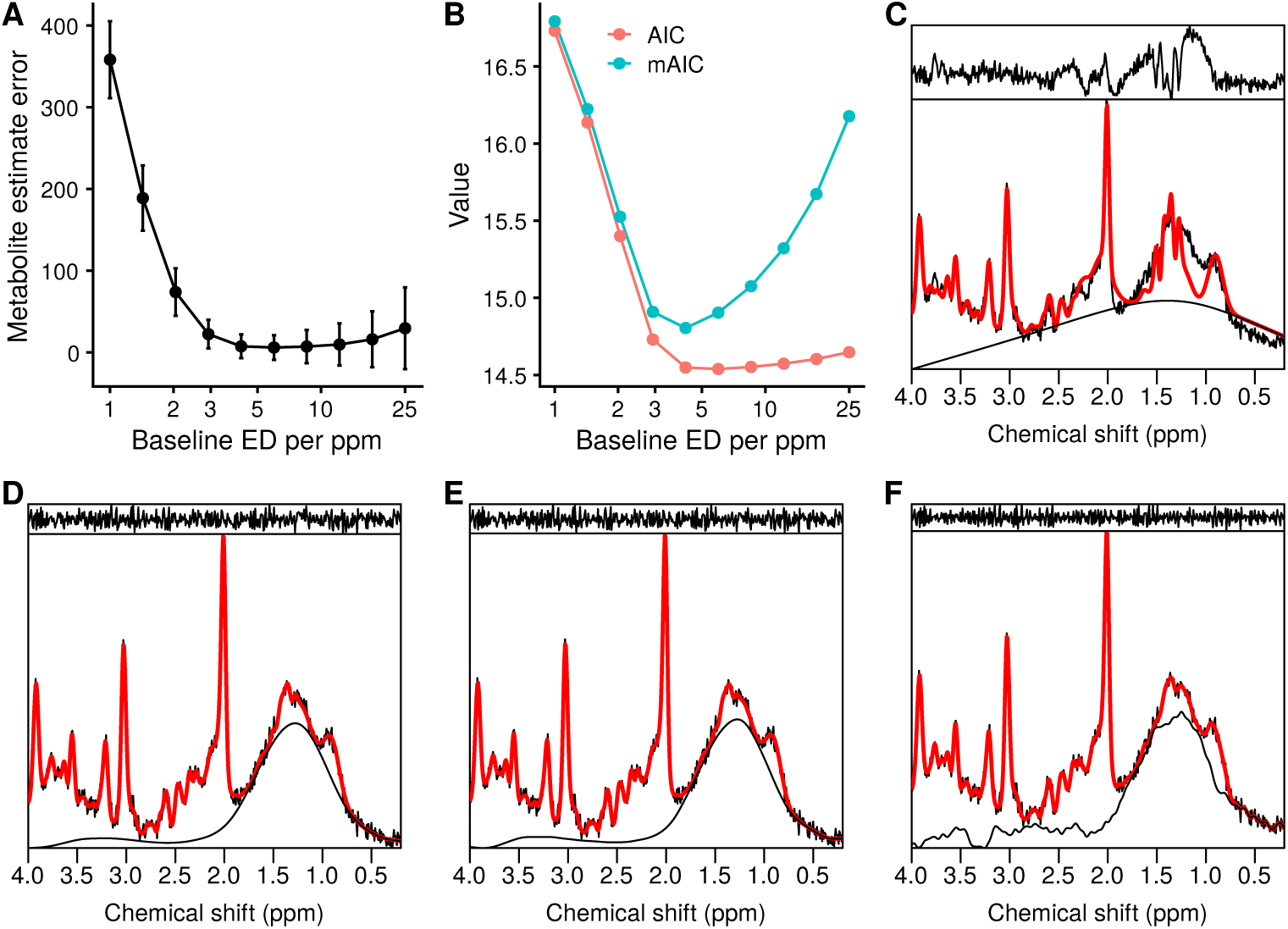
P-spline MRS baseline estimation of simulated data with a broad artifact at 1.3 ppm. A) metabolite estimate error as a function of baseline flexibility. Error bars represent standard deviations, increased by a factor of 5 to aid visualization. B) AIC and modified-AIC as a function of baseline flexibility. C) - F) Fit results with the baseline shown (black) underlying the simulated data (black) and fit (red). The fit residual is shown above the data and fit results are presented for the following values of baseline ED per ppm: C) 1, D) 4.2, E) 6.0, F) 25.

From the results presented in following sections, it was found that the AIC had a tendency to slightly overestimate the optimal level of baseline flexibility, and that a simple modification to Equation 6 improved accuracy:

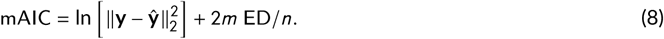

Setting *m* to a value of 5 was empirically found to be a good compromise between bias and variance for all simulated and in vivo analyses presented herein. The value was kept constant for all analyses presented, placing a greater penalty on overly flexible baseline models — resulting in a smoother baseline estimate relative to the standard AIC (Figure 3B).

## 3 METHODS

### 3.1 Adaptive Baseline Fitting Algorithm

In this section we describe a fully-automated ^1^H MRS analysis method based on the P-spline fitting approach presented in the Theory section. The emphasis of the design is to automatically adapt the baseline flexibility for improved accuracy, and we refer to the full algorithm as: *Adaptive Baseline fitting* or ABfit. The algorithm consists of 4 main steps, which will be described in order of execution:

1. Coarse frequency alignment.
2. Approximate iterative fitting.
3. Baseline smoothness estimation.
4. Detailed iterative fitting.

#### 3.1.1 Coarse frequency alignment (step 1)

Unprocessed MRS data typically have an unknown and erroneous frequency offset *f*_*o*_ that displaces all resonances equally. The first step of ABfit is to estimate *f*_*o*_ (measured in Hz) to ensure the basis set of known signals are approximately matched to the acquired data. Raw MRS data is digitally sampled at a frequency of *fs* Hz in the time-domain, and defined a vector of *N* complex data points:

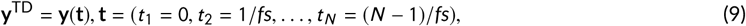

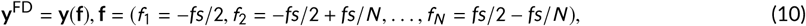

where superscript TD and FD denote time and frequency-domain signal representations — calculated with the discrete Fourier transform. We simulate a reference data set **r**^TD^, with twice the number of data points 2*N* sampled at *fs*, containing three resonances corresponding to the main singlet metabolites typically present in ^1^H MRS data at 2.01, 3.03 and 3.22 ppm. Three resonances are used to ensure the method is appropriate for clinical MRS data, where NAA levels may be very low or absent. The acquired data **y**^TD^ are zero-filled to twice the original length to increase spectral resolution — before convolving with **r**^TD^ to estimate the frequency offset *f*_*o*_ [8].

#### 3.1.2 Approximate iterative fitting (step 2)

The goal of this stage of the algorithm is to find estimates of the three parameters with the largest influence on the fit residual: 1) the signal phase *ϕ*_0_; 2) an approximate lineshape parameter *d*_*g*_ and 3) an improved estimate of *f*_*o*_. The phase and frequency offset are applied to the acquired data as follows:

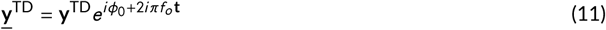

The lineshape parameter applies Gaussian line-broadening to each signal in the basis set **M**, and the scaling factor *β* is introduced to define *d*_*g*_ as the Full Width at Half Maximum (FWHM) measured in Hz:

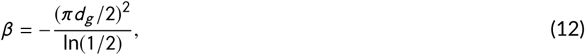

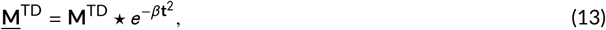

where * denotes element-wise multiplication of the line-broadening function for each column of the basis matrix. A simple, one parameter, lineshape model was chosen for this step to reduce the chance of poor solutions being found by the optimizer due to presence of local minima. Gaussian lineshape broadening was applied to all signals in the basis set **M**, including macromolecues. Metabolite signals in this paper were simulated with pure Lorentzian lineshape to model T2 relaxation — resulting in a Voigt lineshape following the Gaussian broadening applied in this step [31]. Combining the modified basis with the P-spline basis, and solving for **â** in the least-squares sense with the same constraints as the previous section:

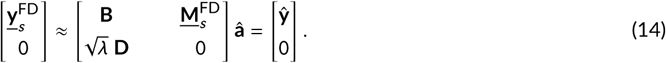

Note that following parametric modification in the time-domain, the data and basis of known signals are zero-filled to twice their original length before being transformed to the frequency-domain. Only the real valued data points in frequency domain between 1.8 and 4 ppm are fit in this stage of the algorithm (denoted by subscript *s*) to reduce the influence of any contaminating lipid signals around 1.3 ppm. The parameters: *ϕ*_0_, *d*_*g*_ and *f*_*o*_ are estimated using Nelder-Mead simplex algorithm with bound constraints [32], minimizing:

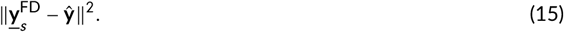

A *λ* value corresponding to 1 ED per ppm is used with a P-spline basis containing 15 spline functions per ppm, resulting in a total of 33 (15 × (4 − 1.8)) components in the basis **B**, with the same density of spline functions being used for all subsequent steps. A value of 1 ED per ppm was empirically found to work well with all data presented herein, however may need to be adjusted for spectra with fundamentally different levels of baseline complexity — such as ^**31**^P MRS. Constraints are placed on the optimization algorithm to restrict the linebroading parameter to be positive and produce additional broadening of less than 15 Hz FWHM. The frequency offset is also constrained to be ±10 Hz different from the coarse frequency alignment (step 1).

#### 3.1.3 Baseline smoothness estimation (step 3)

Following the determination of the frequency offset, spectral phase and approximate lineshape the next step of ABfit is to estimate the optimal value for *λ*. A set of candidate fits are automtically performed with differing level of baseline smoothness, whilst maintaining the three main fitting parameters constant at their optimized values — as determined in the previous step. The maximum candidate baseline flexibility is set to a *λ* value corresponding to 7 ED per ppm, and the minimum value set to an ED of 2 — equivalent to straight line fit. 20 candidate fits are performed with logarithmic intervals between the maximum and minimum values of ED per ppm, where *λ* is back calculated from Equations 4 and 5, and the mAIC is calculated for each fit (Equation 8). The optimal *λ* value, and therefore ED per ppm, is found by selecting the candidate fit with the lowest mAIC. In contrast to the previous step, a wider spectral range of 0.2 to 4 ppm is analyzed to include the majority of conventionally measured metabolites — corresponding to 57 spline functions.

#### 3.1.4 Detailed iterative fitting (step 4)

In the final stage of ABfit, the overall lineshape model is refined alongside *f*_*o*_, *ϕ*_0_ and minor adjustments to the frequencies and linewidths of the known basis signals. An asymmetric lineshape model is generated in the frequency-domain by smoothly adjusting the Gaussian linewidth parameter as a function of frequency — as described by Stancik and Brauns [33]:

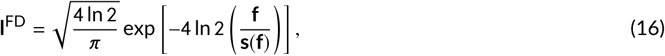

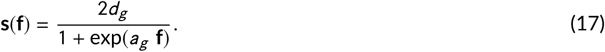

Note, the frequency dependence on the linewidth function is eliminated when the asymmetry parameter *a*_*g*_ = 0, leading to **s** = *d*_*g*_ and pure Gaussian broadening. **l**^FD^ is transformed to the time-domain and applied to each of the basis signals in **M**.

Minor individual changes in the frequency offset and linewidth are applied to each molecular signal in the basis to accommodate variations in the acquired data:

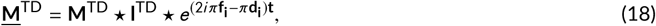

where **f**_**i**_ is a column vector of frequency adjustments measured in Hz, and **d**_**i**_ a column vector of additional Lorenzian line broadening terms measured at FWMH in Hz. Both parameter vectors **f**_**i**_ and **d**_**i**_ are the same length as the number of basis signals in **M**. Each of the global (*f*_*o*_, *ϕ*_0_, *d*_*g*_, *a*_*g*_) and signal specific (**f**_**i**_, **d**_**i**_) parameters are estimated using the Levenberg–Marquardt algorithm [34] implemented with bound constraints, with the same objective function defined in Equations 14 and 15. The optimal baseline smoothness parameter *λ*, (determined in step 3) is used, and the spectral range of 0.2 to 4 ppm included in the iterative optimization procedure. The same parameter constraints were used as in step 2, with the additional lineshape asymmetry parameter *a*_*g*_ being bound between ±0.25, **f**_**i**_ bound between ±1 Hz and 2 ≥ **d**_**i**_ ≥ 0 Hz.

### 3.2 Validation with simulated data

A series of simulations were performed to evaluate the performance of ABfit across a range of common baseline features. For each test, ABfit was performed as described in the previous section — where the baseline flexibility is automatically determined in step 3. Additionally, ABfit was performed with the baseline flexibility set manually (omitting step 3) across a range of 15 values for ED per ppm between 0.53 and 15 to compare with the automated result. Simulated spectra and metabolite estimation errors were calculated as described in section *MRS baseline estimation using P-splines* unless stated otherwise.

An experimentally derived macromolecule profile, comprised of multiple broad resonances combined with fixed relative intensities, was included in the simulations to yield a realistic spectrum [30]. The same macromolecular profile was included in the fitting basis set as a single element for all simulation tests other than fourth and fifth — where a commonly used set of nine individual lipid and macromolecule signals were used in place of the true profile (listed in Table S2). Note the frequencies and linewidths of the main macromolecular resonances are only approximately matched between the experimentally derived macromolecular profile and the set of nine individual lipid and macromolecule signals.

In the first simulation study a broad simulated peak with a Gaussian lineshape (FWHM 100 Hz) at 1.3 ppm was combined with metabolites and the macromolecular signal with a spectral SNR of 54 (Figure 4 and Supporting Information Figure S1). The second simulation study was identical to the first, however the broad simulated peak was removed to yield a perfectly flat baseline (Figures 5 and S2). The third simulation study was identical to the first, however the amplitude of the simulated broad resonance was increased by a factor of 2 (Figures 6 and S3). The data for the fourth simulation study was identical to the second, with a perfectly flat baseline and an experimentally derived macromolecular signal (Figures 7 and S4). However, the experimentally derived macromolecular signal was removed from the fitting basis set and replaced with the set of simulated lipid and macromolecular signals commonly used by default in the LCModel and TARQUIN algorithms (listed in Table S2). The fifth simulation study was identical to the fourth, with the exception of an additional broad simulated peak with a Gaussian lineshape (FWHM 100 Hz) at 1.3 ppm (Figures 8 and S5). The final simulation study was identical to the first, however 12 Hz linebroading was applied to each signal in the basis set (rather than 6 Hz) — resulting in a spectral SNR of 34 (Figures 9 and S6).

**FIGURE 4.**
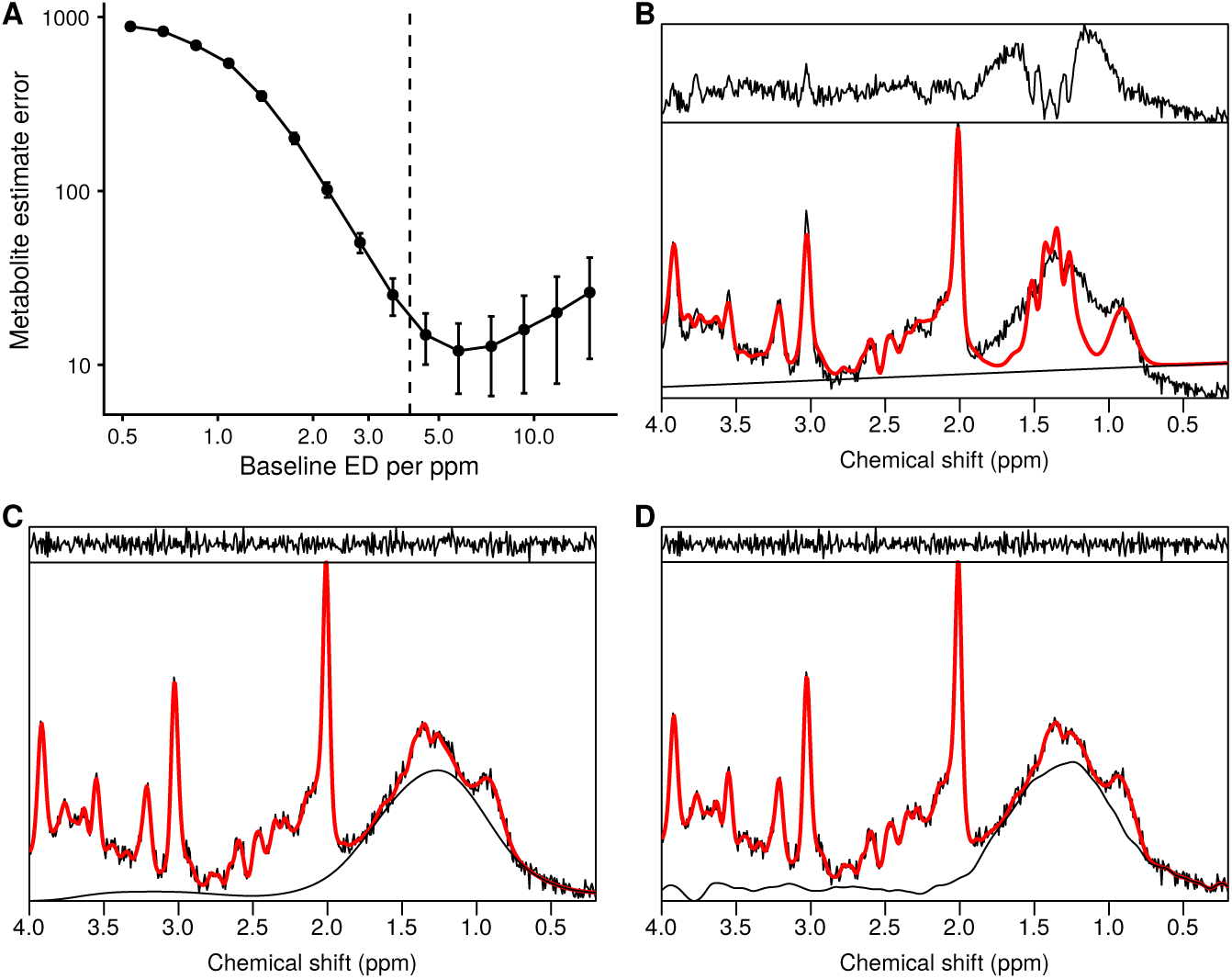
ABfit analysis results for simulated data with a broad baseline distortion at 1.3 ppm. A) metabolite estimate error of ABfit, with the automatically determined level of baseline flexibility (4.1 ED per ppm) shown as a dashed vertical line. Errors values are plotted on a logarithmic scale for clarity. ABfit results with baseline flexibility of B) 0.5, C) 4.1 and D) 15.0 ED per ppm.

**FIGURE 5.**
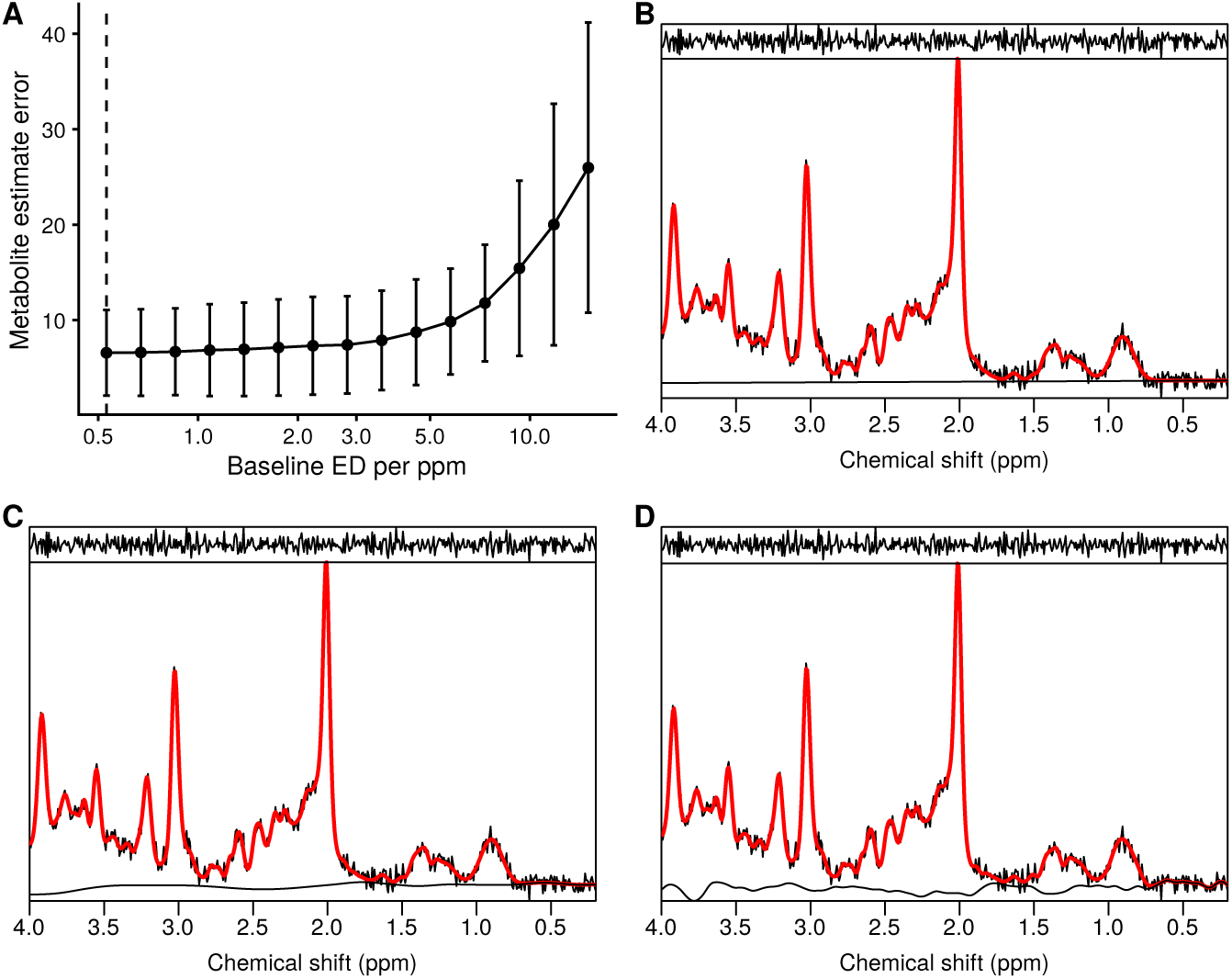
ABfit analysis results for simulated data without baseline distortion. A) metabolite estimate error of ABfit, with the automatically determined level of baseline flexibility (0.53 ED per ppm) shown as a dashed vertical line. ABfit results with baseline flexibility of B) 0.53, C) 4.5 and D) 15.0 ED per ppm.

**FIGURE 6.**
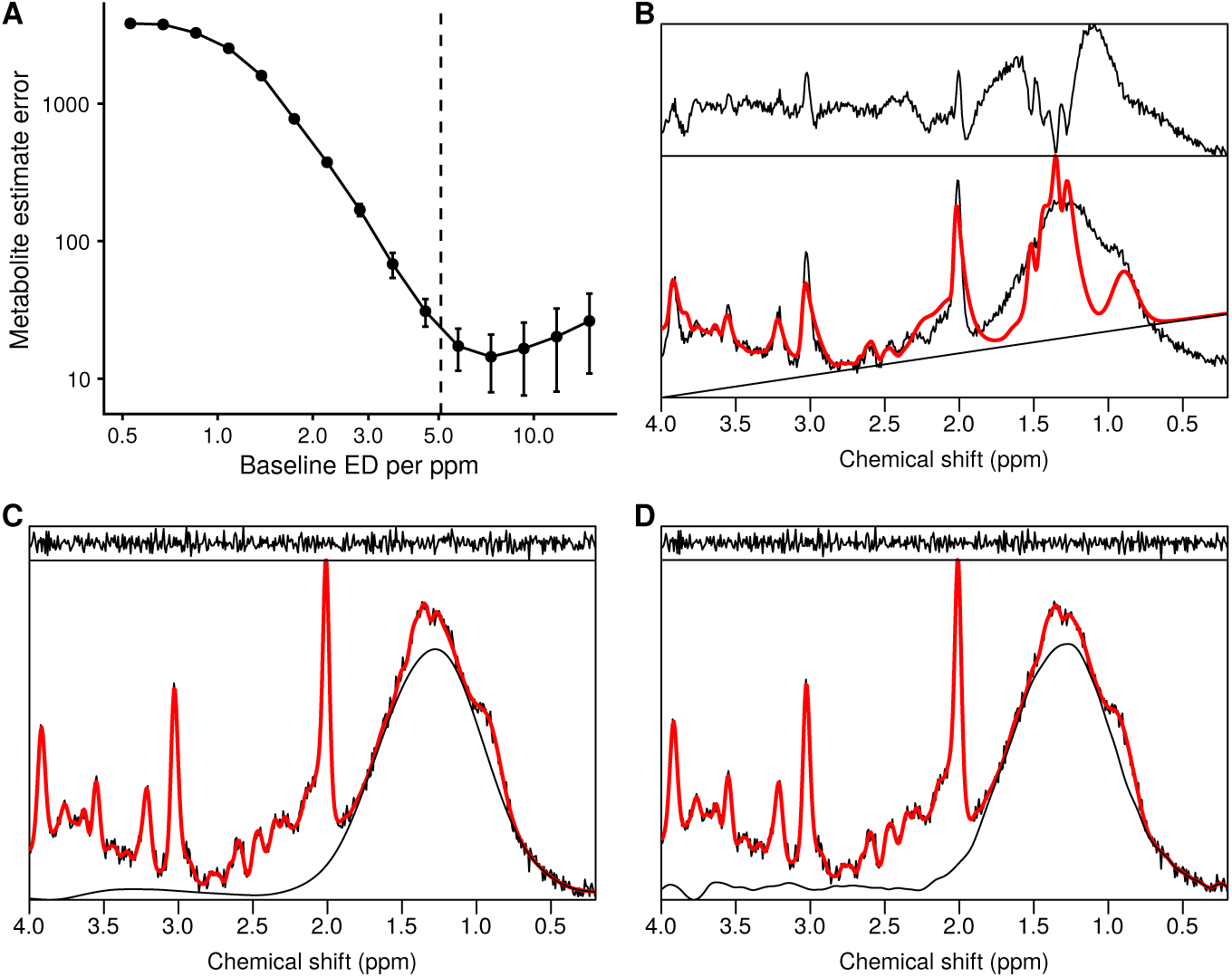
ABfit analysis results for simulated data with a broad baseline distortion at 1.3 ppm with twice the amplitude compared to Figure 4. A) metabolite estimate error of ABfit, with the automatically determined level of baseline flexibility (5.1 ED per ppm) shown as a dashed vertical line. Errors values are plotted on a logarithmic scale for clarity. ABfit results with baseline flexibility of B) 0.5, C) 5.1 and D) 15.0 ED per ppm.

**FIGURE 7.**
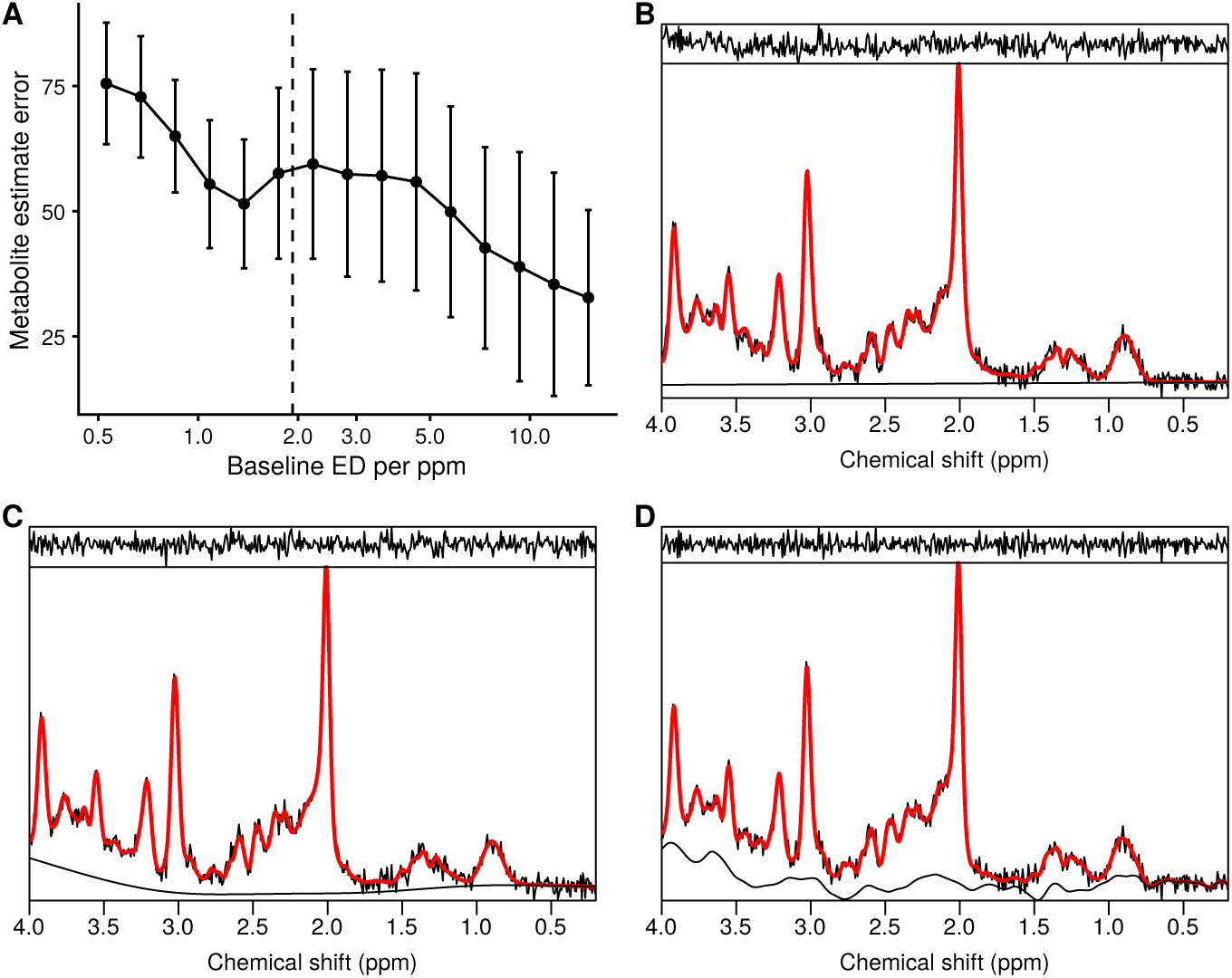
ABfit analysis results for simulated data without baseline distortion, but with the true macromolecular basis signal replaced with individually simulated lipid and macromolecular signals. A) metabolite estimate error of ABfit, with the automatically determined level of baseline flexibility (1.9 ED per ppm) shown as a dashed vertical line. ABfit results with baseline flexibility of B) 0.5, C) 2.0 and D) 15.0 ED per ppm.

**FIGURE 8.**
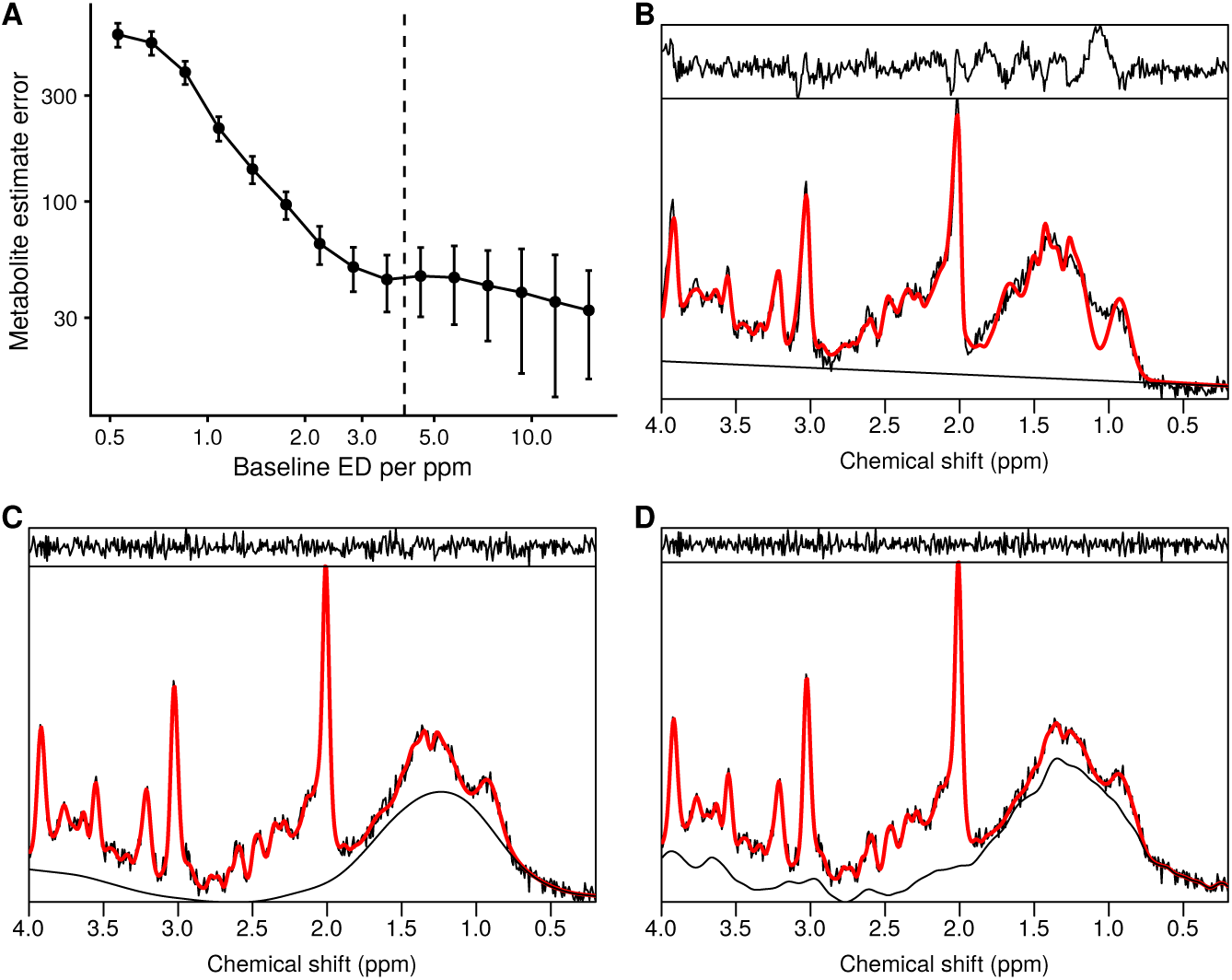
ABfit analysis results for simulated data with a broad baseline distortion at 1.3 ppm and the true macromolecular basis signal replaced with individually simulated lipid and macromolecular signals. A) metabolite estimate error of ABfit, with the automatically determined level of baseline flexibility (4.1 ED per ppm) shown as a dashed vertical line. Errors values are plotted on a logarithmic scale for clarity. ABfit results with baseline flexibility of B) 0.5, C) 4.1 and D) 15.0 ED per ppm.

**FIGURE 9.**
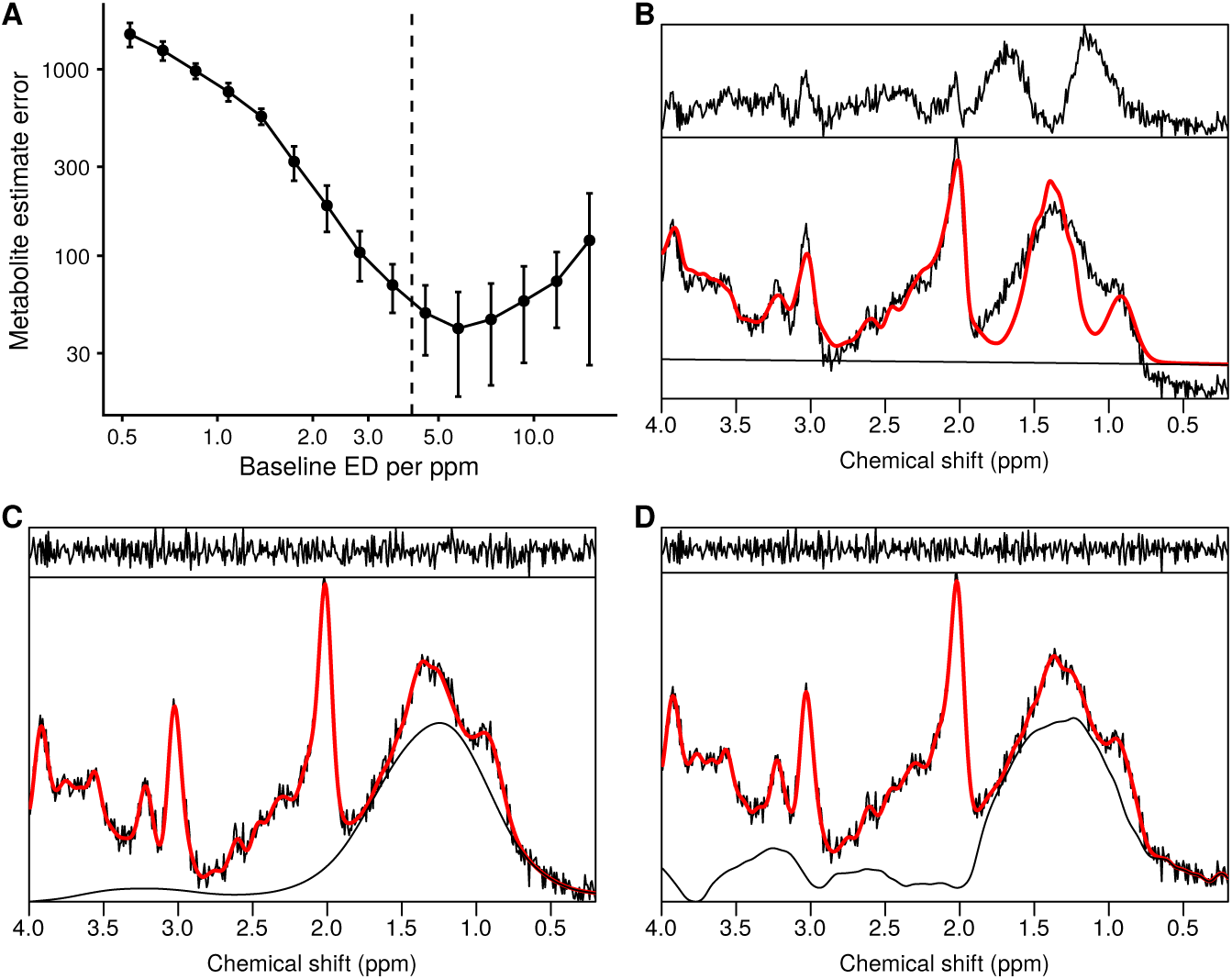
ABfit analysis results for simulated data with a broad baseline distortion at 1.3 ppm and metabolite FWHM of 0.1 ppm. A) metabolite estimate error of ABfit, with the automatically determined level of baseline flexibility (4.1 ED per ppm) shown as a dashed vertical line. Errors values are plotted on a logarithmic scale for clarity. ABfit results with baseline flexibility of B) 0.5, C) 4.1 and D) 15.0 ED per ppm.

### 3.3 Validation with experimentally acquired data

Whilst simulation studies are important to assess the true accuracy of a method, it is challenging to adequately model the true range of variation present in MRS data. Therefore, ABfit was tested on experimentally acquired MRSI data to ensure validity and robustness to common artifacts — such as baseline distortions from scalp lipids, shimming variations and minor shifts in metabolite frequency.

MR data were acquired from two healthy adults with a 3 T Siemens Magnetom Prisma (Siemens Healthcare, Erlangen, Germany) system using a 32-channel receiver head coil-array. T1-weighted MRI was acquired with a 3D-MPRAGE sequence: FOV = 208 x 256 x 256 mm, resolution = 1 x 1 x 1 mm, TE/TR = 2 ms/2000 ms, inversion time = 880 ms, flip angle = 8 degrees and GRAPPA acceleration factor = 2. MRSI data was acquired using 2D MRSI: FOV = 160 x 160 x 15 mm, nominal voxel resolution 10 x 10 x 15mm, TE/TR = 40 ms/2000 ms, complex data points = 1024, sampling frequency = 2000 Hz. The MRSI slice was aligned axially in the subcallosal plane with an approximately 1 mm gap from the upper surface of the corpus callosum. The semi-LASER method [35] was used localize a 100 x 100 x 15 mm ROI, central to the FOV, 4 saturation regions were placed around the ROI prescribing a 100 x 100 mm interior, and an addition 4 saturation regions were positioned to intersect the four corners of the semi-LASER ROI to provide additional scalp lipid suppression. The total MRSI acquisition time was 5 minutes and 6 seconds.

Following acquisition, the four corner voxels were excluded from the central 8 by 8 grid due to their close proximity to the diagonal saturation regions, and ABfit was performed on the remaining 60 voxels. ABfit was applied without any manual adjustments, and exactly the same algorithm was used to analyze the simulated and acquired data. The percentage of white matter, gray matter and CSF contribution to each voxel was measured from segmentation of the T1 MRI using the FAST method [36] as implemented in the FSL software package (v6.0.1). The gray matter fraction was calculated as the percentage volume of gray matter divided by the sum of gray and white matter volumes, and compared with metabolite ratios.

The acquisition of human data included in this study was conducted with the approval of an Institutional Ethics Board.

## 4 RESULTS

### 4.1 Simulation

The first simulation test contained a baseline distortion to mimic a spurious signal from scalp lipids, and the results of ABfit analyses are shown in Figures 4 and S1. The metabolite estimation error plot as a function of baseline flexibility shows the same characteristics as the simpler analysis model used in Figure 3A, however the errors are elevated due to the increased number of parameters in the ABfit model — necessary to handle variations common in acquired MRS data, such as phase and linewidth. The automated estimate for the baseline flexibility (dashed vertical line) represents a reasonable trade-off between bias introduced by insufficient flexibility (part B) and instability — evidenced by increasing standard deviation error bars for greater baseline flexibility (part D). A fit corresponding to the automatically determined baseline flexibility is shown in part C, where the true shape of a broad Gaussian peak centered at 1.3 ppm is apparent from the baseline estimate. A comparison between the true and estimated baseline components is shown in Figure S1 part C with the largest errors shown around 1.3 and 4 ppm. These spectral frequencies correspond to the lactate and alanine resonances which have been overestimated (part B) due to their overlap with the strong baseline artifact and noise.

The baseline distortion was removed for the second simulation study to test the ABfit approach for ideal spectra where the basis set alone is sufficient for accurate analysis. The error plot is shown in Figure 5, illustrating the absence of bias due to baseline underfitting. A compromise between bias and variance is unnecessary in the ideal case, as the correctly determined baseline flexibility has the lowest error and variability. Compared to the first simulation study, the metabolite errors are much smaller and may be attributed to noise (Figure S1 vs S2 part B).

In the third simulation test the amplitude of the broad distortion at 1.3 ppm was doubled compared to the first. A reasonable estimate of the optimal baseline flexibility is found using the ABfit method (Figure 6A) with comparable levels of accuracy relative to the reduced amplitude baseline distortion (Figure 4A). Figure S3 parts B and C show the largest metabolite errors for Lactate and Alanine due to their strong overlap with the baseline distortion at 1.3 ppm.

In the fourth simulation test a set of independent broad lipid and macromolecular signals in the basis were used to model the experimentally derived macromolecular profile. Figure 7A shows the lowest mean errors are obtained at higher levels of baseline flexibility (part D) — indicating a stronger bias at lower level of flexibility (parts B, C) compared to the previous simulations. This is likely explained by inadequate modeling of the broad macromolecular components around 3.8 ppm since these are not present in the commonly used individual macromolecular and lipid basis (Table S2). Figure S4 part D shows a much larger discrepancy between the experimentally derived macromolecular profile used to simulate the data “true” and the estimate “est.” compared to the previous simulation results. A combination of metabolites (part B) and baseline contributions (part C) are used to model the discrepancy in the macromolecular signal components — resulting in increased metabolite errors.

The fifth simulation test was the same as the fourth, except a broad baseline distortion was added at 1.3 ppm. Figure 8 shows a baseline flexibility of 4.1 ED per ppm was necessary to model the baseline distortion at 1.3 ppm and the macromolecular components at 3.8 ppm not present in the basis set. Figure S5 part D shows a greater level of modeling error for the macromolecular components around 1.3 ppm when compared to the previous simulation test, due to their overlap with the baseline distortion.

In the final simulation test, the first test was repeated with broader metabolite signals to evaluate the efficacy of the automated baseline determination for poorly shimmed data. Errors are larger compared to the first test, due to the reduction in SNR and spectral resolution, however a reasonable estimate of the optimal baseline flexibility of approximately 4 ED per ppm is found (Figure 9A). The primary source of metabolite error is shown in Figure S6 part B, where the lactate and alanine signals are confused with the baseline distortion to a greater degree compared to the first simulation test.

### 4.2 Experimental

Significant scalp lipid contamination was found in some of the voxels from the first MRSI scan, most likely resulting from subject movement between the T1 anatomical and MRSI acquisition. ABfit result plots are shown in Figure 10 for a row of eight voxels spanning the localization region — showing a high level of baseline distortion (part A) which becomes increasingly reduced for voxels further from the source. The automatically determined level of baseline flexibility correlates well with the severity of the distortion, and no significant spurious signals are present in the fitting residual.

**FIGURE 10.**
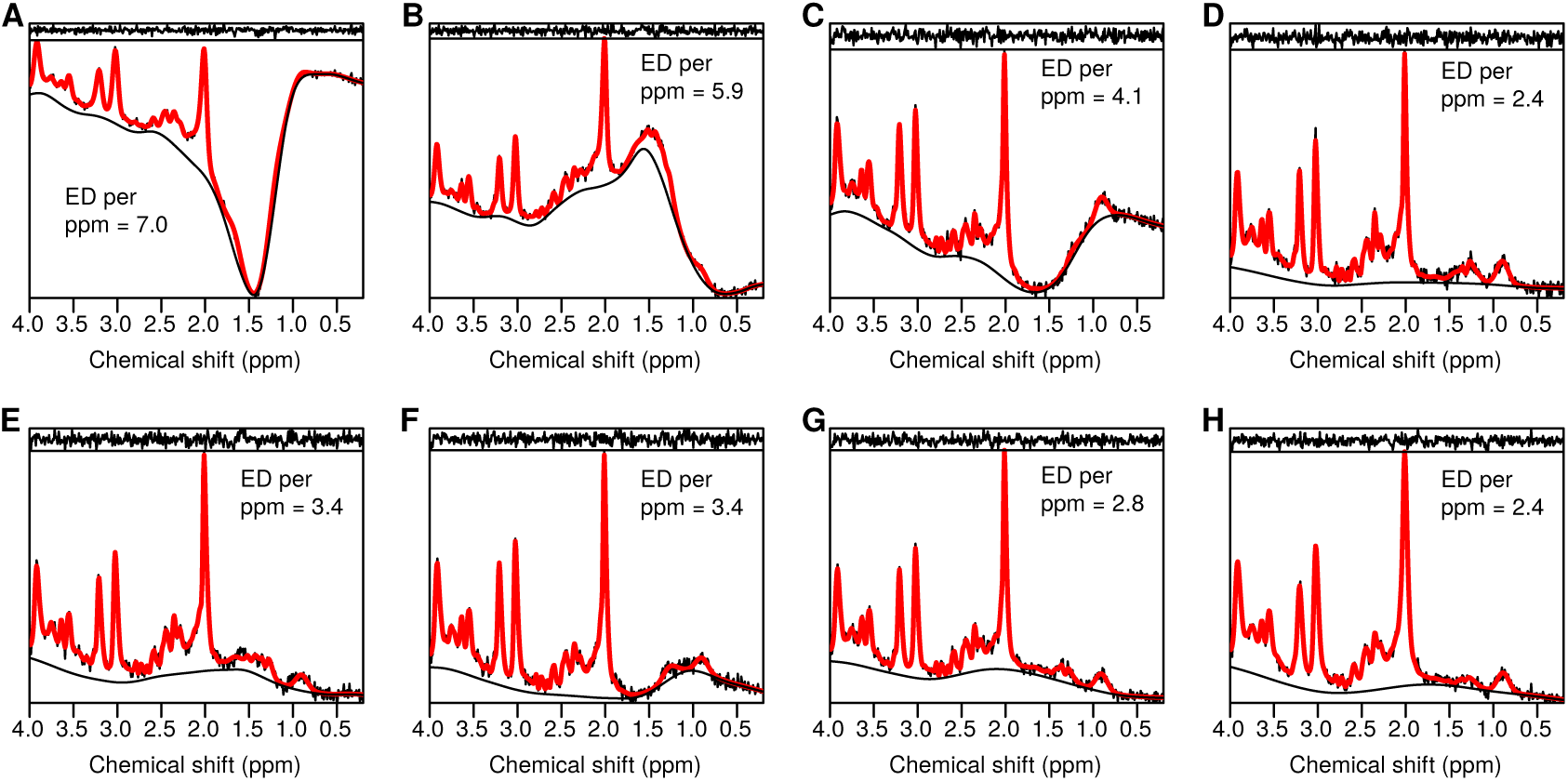
ABfit result plots for a row of 2D MRSI voxels with variable levels of baseline distortion originating from scalp lipid contamination. The automatically determined level of baseline flexibility is listed on each plot.

High quality MRSI was acquired for the second scan with only minimal baseline distortion observed across all analyzed spectra. A strong correlation between metabolite levels and the underlying tissue contribution is visually apparent from Figure S7 part A, with an increased tNAA / tCr ratio in white matter compared to gray matter (tNAA = NAA + NAAG, tCr = PCr + Cr, tCho = GPC + PC, Glx = Glu + Gln). A linear regression of select metabolite ratios with the gray matter fraction is plotted in Figure S7 parts B, C and D, with strong correlations observed — in good agreement with high field observations [37, 38]. The mean full-width at half maximum resolution across the voxels analyzed was 0.032 ppm (3.9 Hz), measured from the tNAA resonance. The mean SNR was 85, with the noise region defined as the real valued data points between -0.5 and -2.5 ppm.

Two example fits from one of the voxels in the second MRSI data set are shown Figure S8. In part A, the ABfit method is applied as described previously, and in part B the lineshape asymmetry parameter (*a*_*g*_) is heavily constrained to enforce a symmetric lineshape model. A smaller fit residual in the tNAA spectral region is found for the asymmetric lineshape model, justifying the minor increase in modeling complexity associated with an additional fit parameter.

## 5 DISCUSSION AND CONCLUSIONS

A new algorithm to automatically determine the optimal level of baseline flexibility has been developed and validated using simulated and acquired MRS data. LCModel is currently the most widely used approach for automatically estimating the optimal baseline flexibility, where the optimal penalty parameter (*α*_*B*_) is chosen by gradually increasing its value until the boundary of the 50% confidence region for the fit is achieved — estimated by comparing successive fits to the first in the series [39, 8]. In contrast to LCModel, the method presented here uses a modification to the AIC to determine the optimal penalty factor — as part of a four-step fitting procedure. An additional difference is the use of P-splines in ABfit compared to smoothing spline approach [40] used in LCModel.

Sima and Van Huffel proposed the use of the classical generalized cross-validation (GCV) criterion, combined with a golden-section search, to determine the optimal penalty parameter value [41]. High accuracy was demonstrated for simulated data, however the method was only tested with good starting values for the non-linear fitting parameters, which are not typically available for experimentally acquired MRS. In ABfit, these non-linear parameters are estimated using a simplified initial fit (step 2) and subsequently refined (step 4). More recently, Zhang and Shen showed that a measure of the baseline uncertainty is also a useful criterion to determine baseline smoothness for simulated and experimentally acquired data [42].

ABfit was shown to find a reasonable compromise between bias and variance for the majority of simulation tests. However, in the fourth simulation test, where a set of independent approximate macromolecular signals were used to fit a realistic macromolecular model, the optimal flexibility was less easily determined. In this case, the AIC penalty for an overly rigid baseline was minor, since a low residual could still be achieved through the increased freedom afforded by using a set of independent macromolecular signals (Figure 7, part B). However, in this case, a more rigid baseline introduces a greater level of metabolite estimation bias due to the mismatch between the simulated and modeled macromolcular profiles. This represents a significant challenge for automated baseline selection for data with lower SNR — a problem previously identified by Near et al [19] using the LCModel package.

Potential solutions to reduce metabolite estimation bias associated with low SNR include the use of more accurate macromolecular modeling [30] and opting to use a fixed level of baseline flexibility. Whist the default approach for ABfit is to automatically select the level of baseline flexibility, fitting options are implemented to specify a fixed degree. Alternatively, baseline flexibility may be systematically adjusted by changing the mAIC scaling parameter *m*, and this may be necessary for data with significantly different baseline characteristics such as ^**31**^P MRS. Each of these approaches has advantages and weaknesses and should be justified depending on the study aims. For example, in functional-MRS a small change in metabolite levels is generally sought, and therefore a less flexible baseline may be preferred — since any metabolite estimation bias is eliminated when the change is normalized to a well determined signal. As a general rule, the baseline should be sufficiently flexible to eliminate any broad features present in the residual. However, poor spectral SNR will mask these features, and therefore caution is advised when comparing metabolite levels between spectra with greatly differing SNR — particularly when macromolecular signals are only partially modeled by the basis set.

In conclusion, new MRS analysis method with adaptive baseline modeling is presented and validated on simulated and experimentally acquired data. The approach is fully-automated and integrated into a free and open-source software package — providing a transparent and reproducible platform for future MRS studies [43].

## DATA AVAILABILITY STATEMENT

The code and data that supports the findings of this study are openly available in Github at https://github.com/martin3141/abfit_paper, tag “final” and https://github.com/martin3141/spant, tag “v1.4.0”.

## REFERENCES

[1] Oz G, Alger JR, Barker PB, Bartha R, Bizzi A, Boesch C, et al. Clinical proton MR spectroscopy in central nervous system disorders. Radiology 2014;270(3):658–79.

[2] Lally PJ, Montaldo P, Oliveira V, Soe A, Swamy R, Bassett P, et al. Magnetic resonance spectroscopy assessment of brain injury after moderate hypothermia in neonatal encephalopathy: a prospective multicentre cohort study. Lancet Neurol 2019;18(1):35–45.

[3] Merritt K, Egerton A, Kempton MJ, Taylor MJ, McGuire PK. Nature of glutamate alterations in schizophrenia: a meta-analysis of proton magnetic resonance spectroscopy studies. JAMA Psychiat 2016;73(7):665–74.

[4] Jelen LA, King S, Mullins PG, Stone JM. Beyond static measures: A review of functional magnetic resonance spectroscopy and its potential to investigate dynamic glutamatergic abnormalities in schizophrenia. J Psychopharmacol 2018;32(5):497–508.

[5] Chen C, Sigurdsson HP, Pepes SE, Auer DP, Morris PG, Morgan PS, et al. Activation induced changes in GABA: Functional MRS at 7T with MEGA-sLASER. Neuroimage 2017;156:207–213.

[6] Wilson M, Andronesi O, Barker PB, Bartha R, Bizzi A, Bolan PJ, et al. Methodological consensus on clinical proton MRS of the brain: Review and recommendations. Magn Reson Med 2019;82(2):527–550.

[7] Cudalbu C, Mlynarik V, Gruetter R. Handling macromolecule signals in the quantification of the neurochemical profile. J Alzheimers Dis 2012;31 Suppl 3:S101–15.

[8] Provencher SW. Estimation of metabolite concentrations from localized in vivo proton NMR spectra. Magn Reson Med 1993;30(6):672–9.

[9] Poullet JB, Sima DM, Simonetti AW, De Neuter B, Vanhamme L, Lemmerling P, et al. An automated quantitation of short echo time MRS spectra in an open source software environment: AQSES. NMR Biomed 2007;20(5):493–504.

[10] Ratiney H, Sdika M, Coenradie Y, Cavassila S, van Ormondt D, Graveron-Demilly D. Time-Domain Semi-Parametric Estimation Based on a Metabolite Basis Set. NMR Biomed 2005;18(1):1–13.

[11] Wilson M, Reynolds G, Kauppinen RA, Arvanitis TN, Peet AC. A constrained least-squares approach to the automated quantitation of in vivo 1 H magnetic resonance spectroscopy data. Magn Reson Med 2011;65(1):1–12.

[12] Young K, Soher BJ, Maudsley AA. Automated spectral analysis II: application of wavelet shrinkage for characterization of non-parameterized signals. Magn Reson Med 1998;40(6):816–21.

[13] Seeger U, Klose U, Mader I, Grodd W, Nägele T. Parameterized evaluation of macromolecules and lipids in proton MR spectroscopy of brain diseases. Magn Reson Med 2003 Jan49:19–28.

[14] Pfeuffer J, Tkac I, Provencher SW, Gruetter R. Toward an in vivo neurochemical profile: quantification of 18 metabolites in short-echo-time (1)H NMR spectra of the rat brain. J Magn Reson 1999;141(1):104–20.

[15] Deelchand DK, Marjanska M, Hodges JS, Terpstra M. Sensitivity and specificity of human brain glutathione concentrations measured using short-TE (1)H MRS at 7 T. NMR Biomed 2016;29(5):600–6.

[16] Terpstra M, Ugurbil K, Tkac I. Noninvasive quantification of human brain ascorbate concentration using 1H NMR spectroscopy at 7 T. NMR Biomed 2010;23(3):227–32.

[17] Marjanska M, Deelchand DK, Hodges JS, McCarten JR, Hemmy LS, Grant A, et al. Altered macromolecular pattern and content in the aging human brain. NMR Biomed 2018;31(2):e3865.

[18] Ratiney H, Coenradie Y, Cavassila S, van Ormondt D, Graveron-Demilly D. Time-domain quantitation of 1H short echotime signals: background accommodation. Mag Reson Mater Phy 2004;16(6):284–96.

[19] Near J, Andersson J, Maron E, Mekle R, Gruetter R, Cowen P, et al. Unedited in vivo detection and quantification of gamma-aminobutyric acid in the occipital cortex using short-TE MRS at 3 T. NMR Biomed 2013;26(11):1353–62.

[20] Giapitzakis IA, Borbath T, Murali-Manohar S, Avdievich N, Henning A. Investigation of the influence of macromolecules and spline baseline in the fitting model of human brain spectra at 9.4 T. Magn Reson Med 2019;81(2):746–758.

[21] Wenger KJ, Hattingen E, Harter PN, Richter C, Franz K, Steinbach JP, et al. Fitting algorithms and baseline correction influence the results of non-invasive in vivo quantitation of 2-hydroxyglutarate with (1)H-MRS. NMR Biomed 2019;32(1):e4027.

[22] Ruppert D, Wand MP, Carroll RJ. Semiparametric Regression. Cambridge Series in Statistical and Probabilistic Mathematics, Cambridge University Press; 2003.

[23] De Boor C. A practical guide to splines; rev. ed. Applied mathematical sciences, Berlin: Springer; 2001.

[24] Eilers PHC, Marx BD. Flexible smoothing with B-splines and penalties. Stat Sci 1996 05;11(2):89–121.

[25] Hastie T, Tibshirani R. Generalized Additive Models. London: Chapman and Hall; 1990.

[26] Akaike H. Maximum likelihood identification of Gaussian autoregressive moving average models. Biometrika 1973 08;60(2):255–265.

[27] Govind V, Young K, Maudsley AA. Corrigendum: proton NMR chemical shifts and coupling constants for brain metabolites. NMR Biomed 2015 Jul;28(7):923–924.

[28] Lawson CL, Hanson RJ. Solving least squares problems, vol. 15 of Classics in Applied Mathematics. Philadelphia, PA: Society for Industrial and Applied Mathematics (SIAM); 1995.

[29] de Graaf RA. In Vivo NMR Spectroscopy: Principles and Techniques. John Wiley & Sons, Ltd.; 2018.

[30] Birch R, Peet AC, Dehghani H, Wilson M. Influence of macromolecule baseline on (1)H MR spectroscopic imaging reproducibility. Magn Reson Med 2017;77(1):34–43.

[31] Marshall I, Higinbotham J, Bruce S, Freise A. Use of Voigt lineshape for quantification of in vivo 1H spectra. Magn Reson Med 1997 May;37:651–657.

[32] Box MJ. A mew method of constrained optimization and a comparison with other methods. Comput J 1965 04;8(1):42–52.

[33] Stancik AL, Brauns EB. A simple asymmetric lineshape for fitting infrared absorption spectra. Vib Spectrosc 2008;47(1):66–69.

[34] Levenberg K. A method for the solution of certain non-linear problems in least squares. Q Appl Math 1944;2(2):164–168.

[35] Scheenen TWJ, Klomp DWJ, Wijnen JP, Heerschap A. Short echo time 1H-MRSI of the human brain at 3T with minimal chemical shift displacement errors using adiabatic refocusing pulses. Magn Reson Med 2008;59(1):1–6.

[36] Zhang Y, Brady M, Smith S. Segmentation of brain MR images through a hidden Markov random field model and the expectation-maximization algorithm. IEEE Trans Med Imag 2001;20(1):45–57.

[37] Nassirpour S, Chang P, Henning A. High and ultra-high resolution metabolite mapping of the human brain using (1)H FID MRSI at 9.4T. NeuroImage 2018;168:211–221.

[38] Hangel G, Strasser B, Povazan M, Heckova E, Hingerl L, Boubela R, et al. Ultra-high resolution brain metabolite mapping at 7 T by short-TR Hadamard-encoded FID-MRSI. NeuroImage 2018;168:199–210.

[39] Provencher SW. A constrained regularization method for inverting data represented by linear algebraic or integral equations. Comput Phys Commun 1982;27(3):213–227.

[40] O’Sullivan F. A Statistical Perspective on Ill-Posed Inverse Problems. Stat Sci 1986 11;1(4):502–518.

[41] Sima DM, Huffel SV. Regularized semiparametric model identification with application to nuclear magnetic resonance signal quantification with unknown macromolecular base-line. J Roy Stat Soc B 2006;68(3):383–409.

[42] Zhang Y, Shen J. Smoothness of in vivo spectral baseline determined by mean-square error. Magn Reson Med 2014;72(4):913–22.

[43] Stikov N, Trzasko JD, Bernstein MA. Reproducibility and the future of MRI research. Magn Reson Med 2019 Dec;82(6):1981–1983.

