## Supporting Information for "Adaptive Baseline Fitting for ^1^H MR Spectroscopy Analysis"

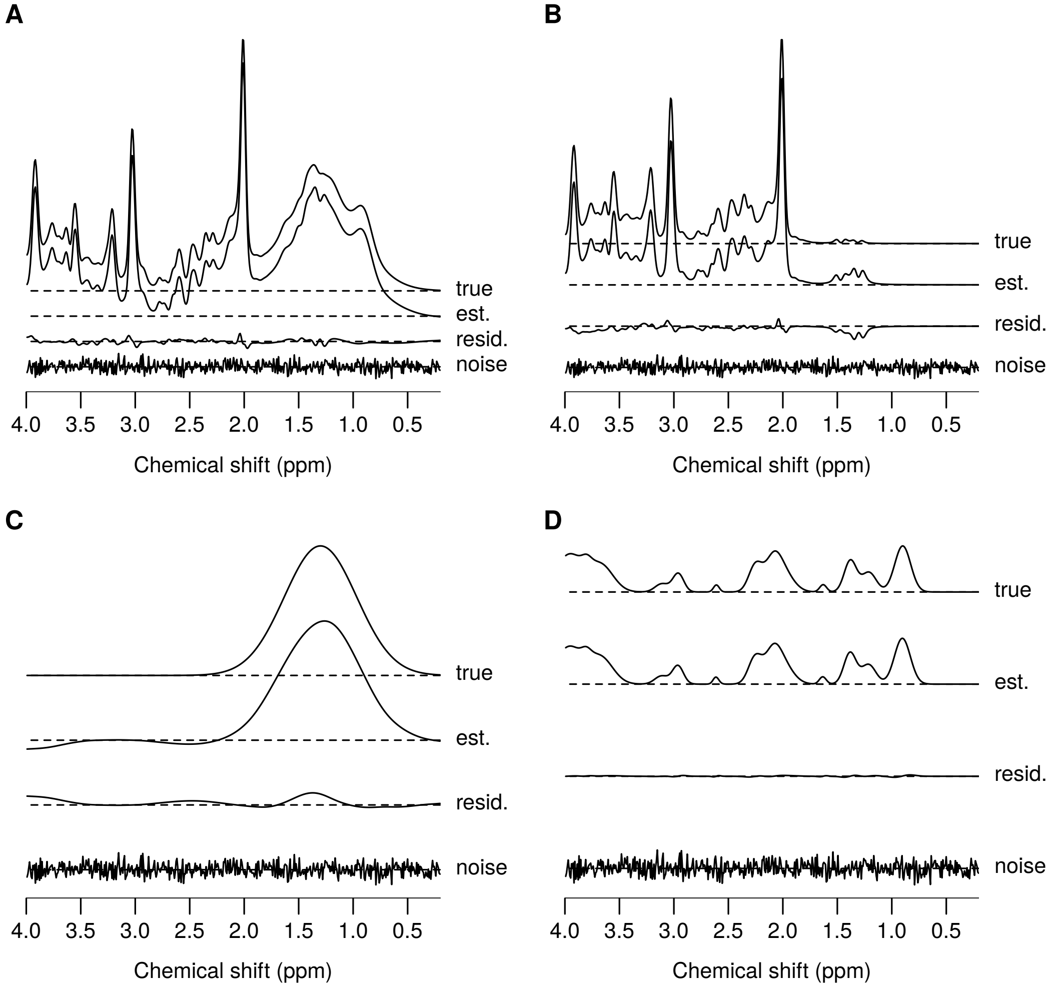


**FIGURE S1** Example ABfit analysis result for simulated data with a broad baseline distortion at 1.3 ppm. Fitting was performed on data comprised of known metabolite, baseline, macromoleclar and noise components (part A "true" + "noise"). Parts B, C and D compare the true and estimated signals separately for the metabolite, baseline and macromolecular components respectively. The simulated noise-free signal ("true") is shown in each subplot for comparison with the estimate from ABfit ("est."). The difference between the true and estimated ("true" - "est.") signals are also shown ("resid"). Horizontal dashed lines represent an intensity of zero for each of the four traces.

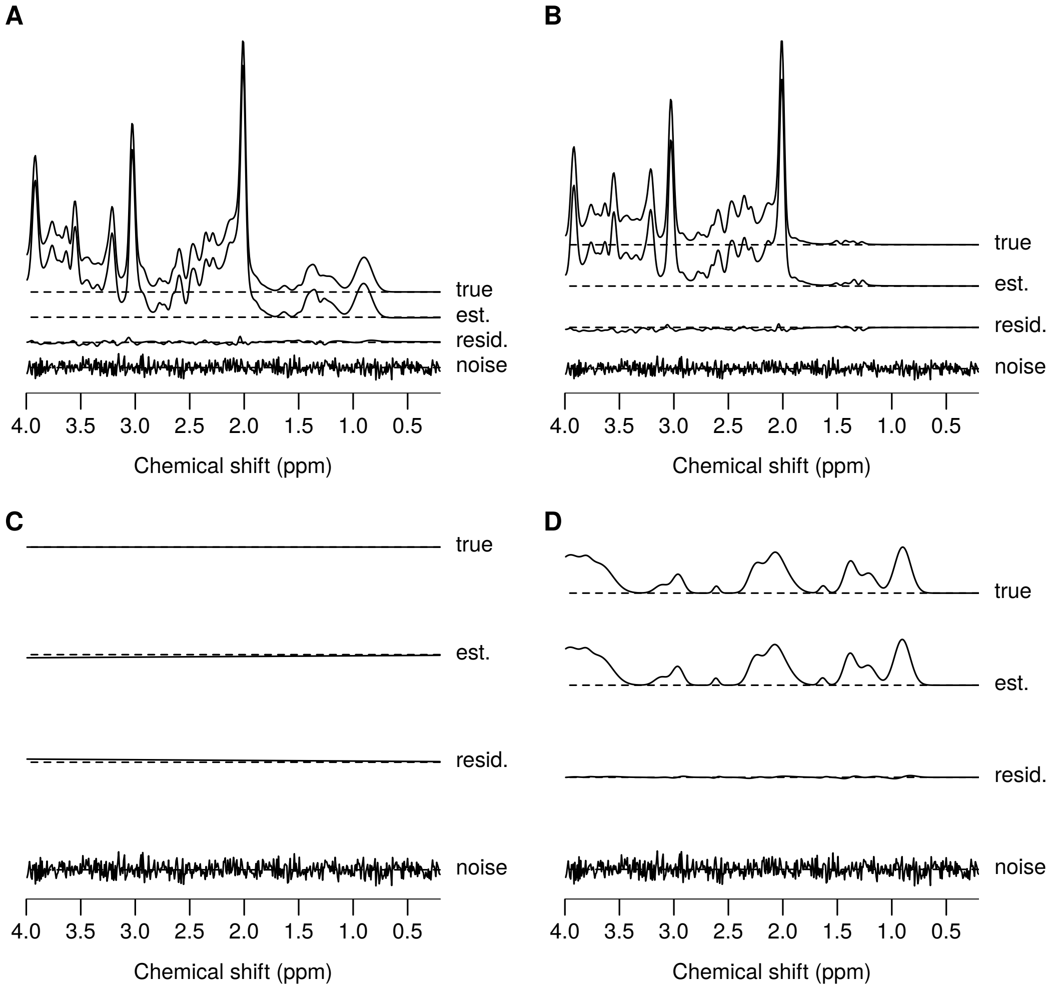


**FIGURE S2** Example ABfit analysis result for simulated data without baseline distortion. Fitting was performed on data comprised of known metabolite, baseline, macromoleclar and noise components (part A "true" + "noise"). Parts B, C and D compare the true and estimated signals separately for the metabolite, baseline and macromolecular components respectively. The simulated noise-free signal ("true") is shown in each subplot for comparison with the estimate from ABfit ("est."). The difference between the true and estimated ("true" - "est.") signals are also shown ("resid"). Horizontal dashed lines represent an intensity of zero for each of the four traces.

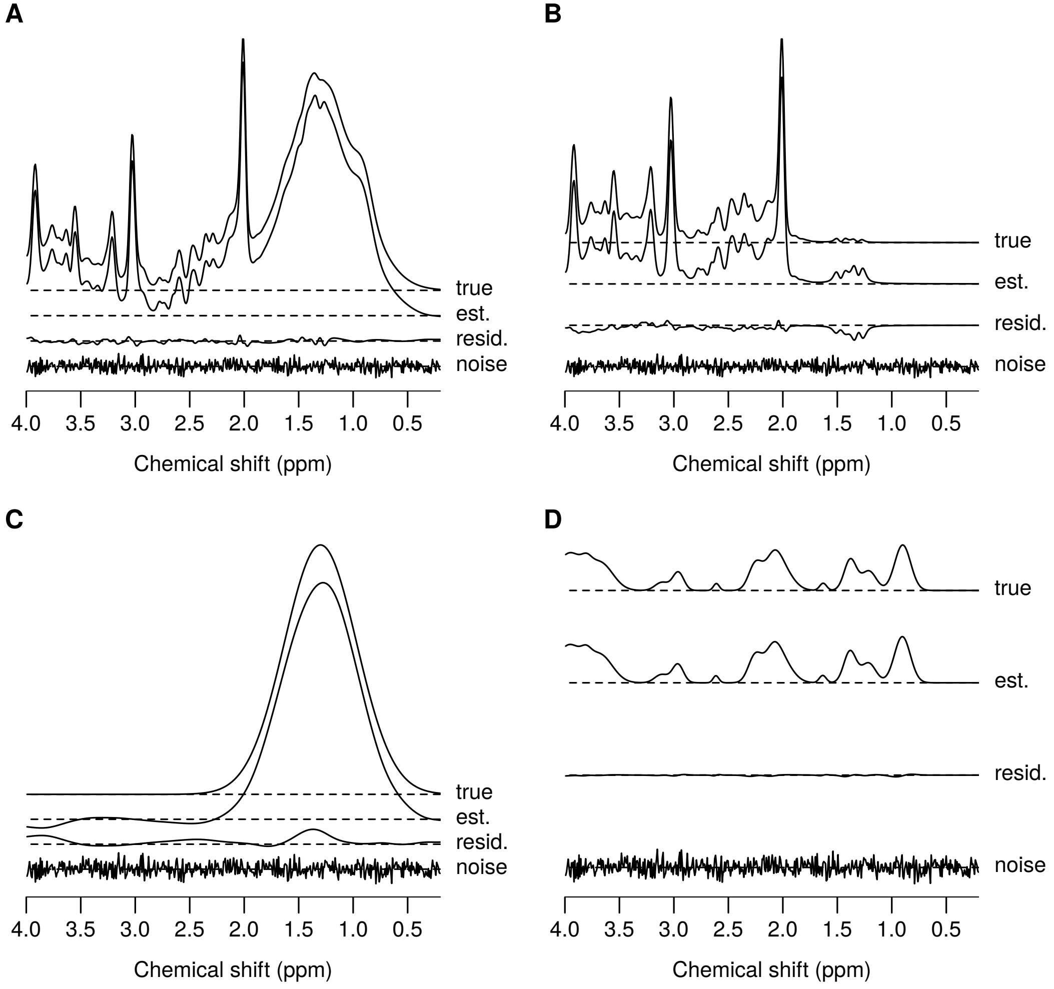


**FIGURE S3** Example ABfit analysis result for simulated data with a broad baseline distortion at 1.3 ppm with twice the amplitude compared to Figures 4 and S1. Fitting was performed on data comprised of known metabolite, baseline, macromoleclar and noise components (part A “true” + “noise”). Parts B, C and D compare the true and estimated signals separately for the metabolite, baseline and macromolecular components respectively. The simulated noise-free signal (“true”) is shown in each subplot for comparison with the estimate from ABfit (“est.”). The difference between the true and estimated (“true” - “est.”) signals are also shown (“resid”). Horizontal dashed lines represent an intensity of zero for each of the four traces.

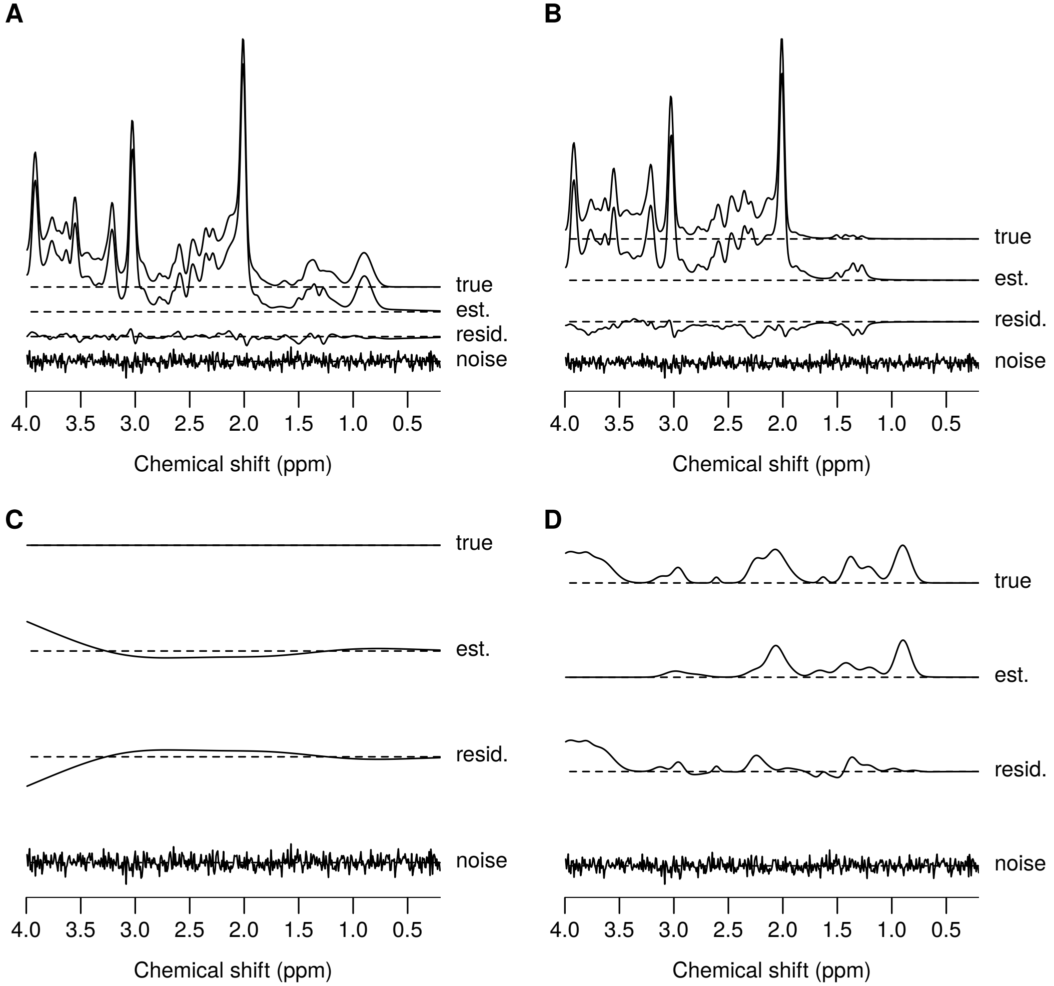


**FIGURE S4** Example ABfit analysis result for simulated data without baseline distortion, but with the true macromolecular basis signal replaced with individually simulated lipid and macromolecular signals. Fitting was performed on data comprised of known metabolite, baseline, macromoleclar and noise components (part A “true” + “noise”). Parts B, C and D compare the true and estimated signals separately for the metabolite, baseline and macromolecular components respectively. The simulated noise-free signal (“true”) is shown in each subplot for comparison with the estimate from ABfit (“est.”). The difference between the true and estimated (“true” - “est.”) signals are also shown (“resid”). Horizontal dashed lines represent an intensity of zero for each of the four traces.

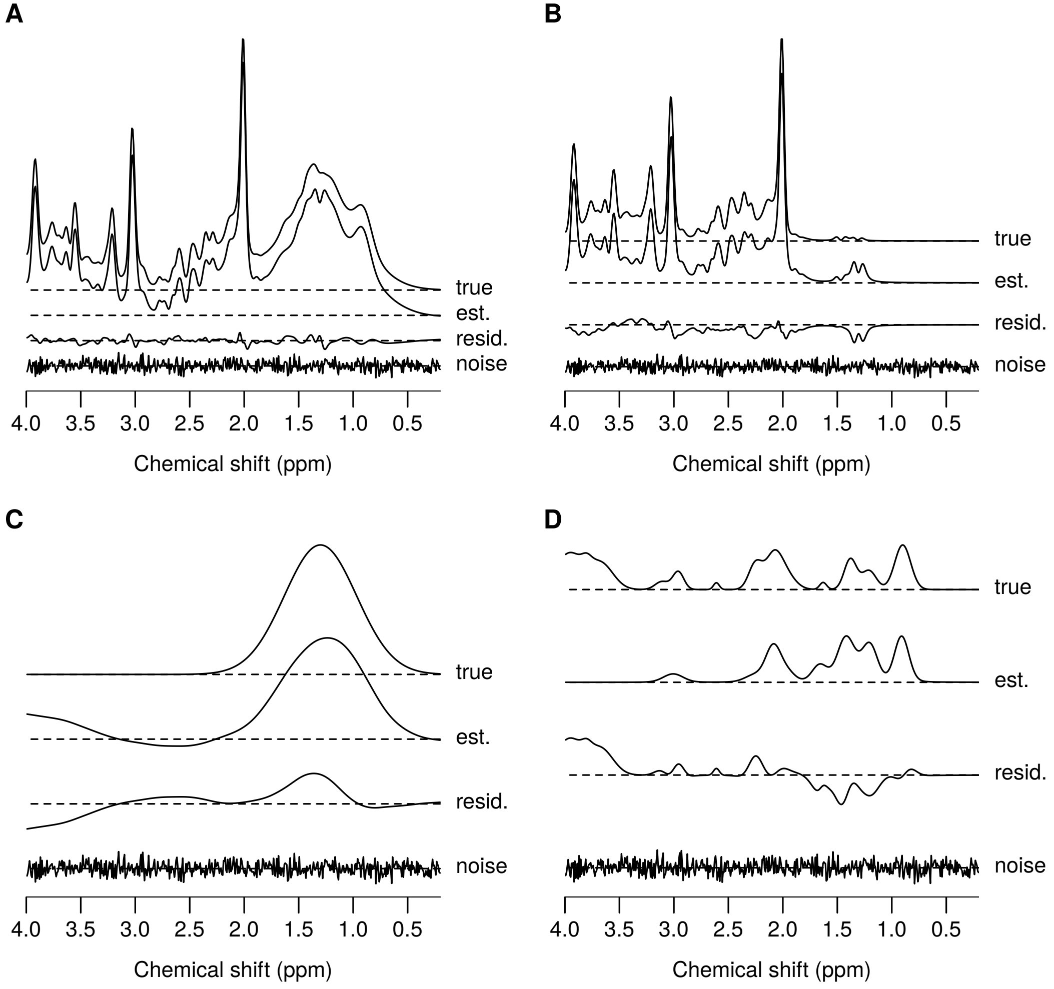


**FIGURE S5** Example ABfit analysis result for simulated data with a broad baseline distortion at 1.3 ppm and the true macromolecular basis signal replaced with individually simulated lipid and macromolecular signals. Fitting was performed on data comprised of known metabolite, baseline, macromoleclar and noise components (part A "true" + "noise"). Parts B, C and D compare the true and estimated signals separately for the metabolite, baseline and macromolecular components respectively. The simulated noise-free signal ("true") is shown in each subplot for comparison with the estimate from ABfit ("est."). The difference between the true and estimated ("true" - "est.") signals are also shown ("resid"). Horizontal dashed lines represent an intensity of zero for each of the four traces.

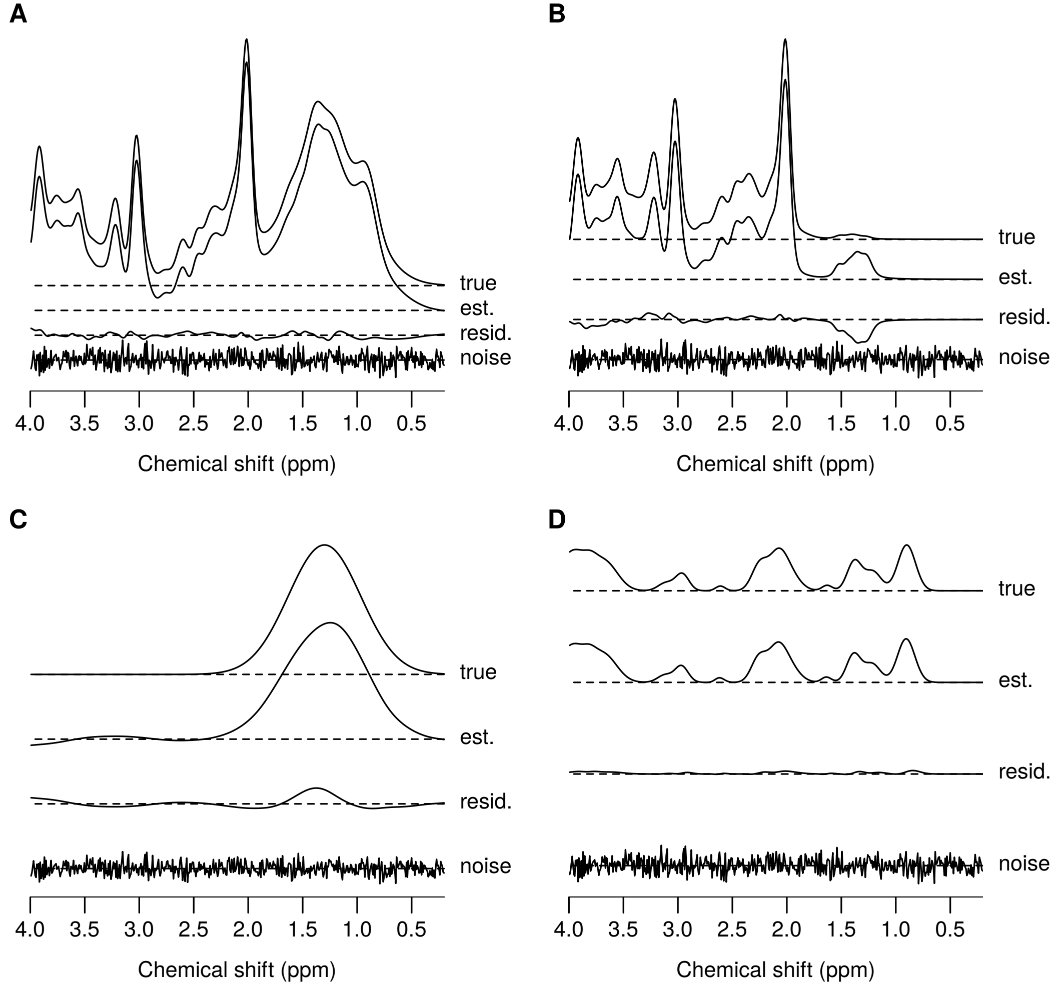


**FIGURE S6** Example ABfit analysis result for simulated data with a broad baseline distortion at 1.3 ppm and metabolite FWHM of 0.1 ppm. Fitting was performed on data comprised of known metabolite, baseline, macromoleclar and noise components (part A “true” + “noise”). Parts B, C and D compare the true and estimated signals separately for the metabolite, baseline and macromolecular components respectively. The simulated noise-free signal (“true”) is shown in each subplot for comparison with the estimate from ABfit (“est.”). The difference between the true and estimated (“true” - “est.”) signals are also shown (“resid”). Horizontal dashed lines represent an intensity of zero for each of the four traces.

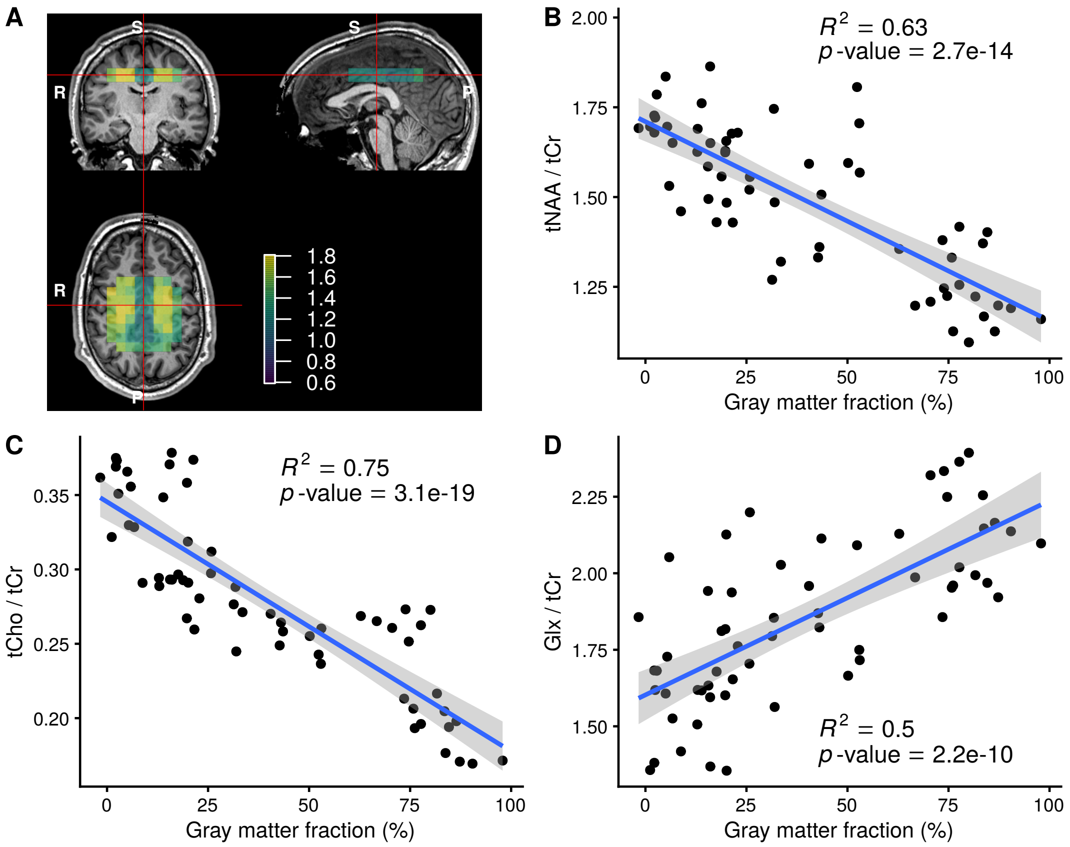


**FIGURE S7** ABfit results from a 2D MRSI semi-LASER acquisition. A) Orthogonal T1 MRI slices intersecting the analysis region — shown as a colored tNAA / tCr metabolite map overlay. Linear regression of key metabolite ratios with the gray matter fraction are shown in parts B, C and D. The line of best fit is plotted in blue with the 95% confidence region in gray.

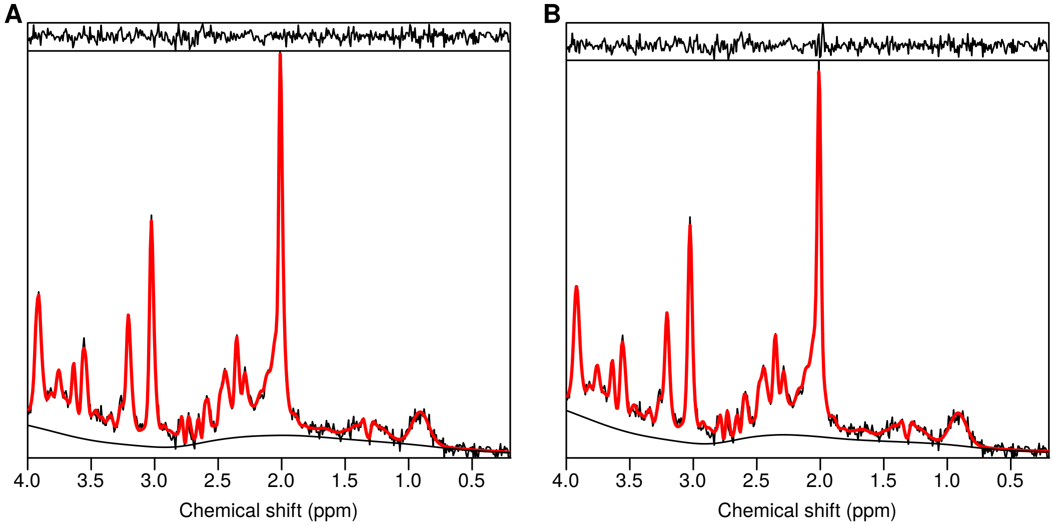


**FIGURE S8** ABfit result plot for a voxel within a 2D MRSI semi-LASER acquisition with an asymmetric lineshape. A) default analysis with an asymmetric lineshape fit model, B) analysis with the lineshape asymmetry parameter ($a_g$) restricted to ±0.0001.

| Metabolite | Amplitude (mM) |
|---|---|
| Alanine | 0.80 |
| Aspartate | 1.00 |
| Creatine | 7.50 |
| gamma-Aminobutyric acid | 1.50 |
| Glucose | 1.50 |
| Glutamine | 4.50 |
| Glutamate | 9.25 |
| Glutathione | 2.25 |
| Glycerophosphorylcholine | 1.00 |
| Lactate | 0.60 |
| myo-Inositol | 6.50 |
| N-acetylaspartate | 12.25 |
| N-acetylaspartylglutamate | 1.50 |
| Phosphocholine | 0.60 |
| Phosphocreatine | 4.25 |
| scyllo-Inositol | 0.35 |
| Taurine | 4.00 |

**TABLE S1** Metabolite concentrations consistent with levels measured in normal brain tissue.

| Signal | Frequency (PPM) | FWHM (PPM) | Amplitude (a.u.) |
|---|---|---|---|
| Lip13a | 1.28 | 0.15 | 2.0 |
| Lip13b | 1.28 | 0.089 | 2.0 |
| Lip09 | 0.89 | 0.14 | 3.0 |
| MM09 | 0.91 | 0.14 | 3.0 |
| Lip20 | 2.04 | 0.15 | 1.33 |
| Lip20 | 2.25 | 0.15 | 0.67 |
| Lip20 | 2.80 | 0.20 | 0.87 |
| MM20 | 2.08 | 0.15 | 1.33 |
| MM20 | 2.25 | 0.20 | 0.33 |
| MM20 | 1.95 | 0.15 | 0.33 |
| MM20 | 3.00 | 0.20 | 0.4 |
| MM12 | 1.21 | 0.15 | 2.0 |
| MM14 | 1.43 | 0.17 | 2.0 |
| MM17 | 1.67 | 0.15 | 2.0 |

**TABLE S2** Parameters used to generate the individual simulated lipid and macromolecule basis signals. Listed components with the same name were summed to form a composite signal.
